# O-GlcNAcylation of FOXK1 orchestrates the E2F pathway and promotes oncogenesis

**DOI:** 10.1101/2024.03.01.582838

**Authors:** Louis Masclef, Oumaima Ahmed, Nicholas Iannantuono, Jessica Gagnon, Mila Gushul-Leclaire, Karine Boulay, Benjamin Estavoyer, Mohamed Echbicheb, Marty Poy, Kalidou Ali Boubacar, Amina Boubekeur, Saad Menggad, Alejandro Schcolnik-Cabrera, Aurelio Balsalobre, Eric Bonneil, Pierre Thibault, Laura Hulea, Yoshiaki Tanaka, Frédérick Antoine-Mallette, Jacques Drouin, El Bachir Affar

**Author notes:** These authors contributed equally to this work (Louis Masclef, Oumaima Ahmed).

## Abstract

Gene transcription is a highly regulated process, and deregulation of transcription factors activity underlies numerous pathologies including cancer. Albeit near four decades of studies have established that the E2F pathway is a core transcriptional network that govern cell division in multi-cellular organisms^1,2^, the molecular mechanisms that underlie the functions of E2F transcription factors remain incompletely understood. FOXK1 and FOXK2 transcription factors have recently emerged as important regulators of cell metabolism, autophagy and cell differentiation^3–6^. While both FOXK1 and FOXK2 interact with the histone H2AK119ub deubiquitinase BAP1 and possess many overlapping functions in normal biology, their specific functions as well as deregulation of their transcriptional activity in cancer is less clear and sometimes contradictory^7–13^. Here, we show that elevated expression of FOXK1, but not FOXK2, in primary normal cells promotes transcription of E2F target genes associated with increased proliferation and delayed entry into cellular senescence. FOXK1 expressing cells are highly prone to cellular transformation revealing important oncogenic properties of FOXK1 in tumor initiation. High expression of FOXK1 in patient tumors is also highly correlated with E2F gene expression. Mechanistically, we demonstrate that FOXK1, but not FOXK2, is specifically modified by O-GlcNAcylation. FOXK1 O-GlcNAcylation is modulated during the cell cycle with the highest levels occurring during the time of E2F pathway activation at G1/S. Moreover, loss of FOXK1 O-GlcNAcylation impairs FOXK1 ability to promote cell proliferation, cellular transformation and tumor growth. Mechanistically, expression of FOXK1 O-GlcNAcylation-defective mutants results in reduced recruitment of BAP1 to gene regulatory regions. This event is associated with a concomitant increase in the levels of histone H2AK119ub and a decrease in the levels of H3K4me1, resulting in a transcriptional repressive chromatin environment. Our results define an essential role of O-GlcNAcylation in modulating the functions of FOXK1 in controlling the cell cycle of normal and cancer cells through orchestration of the E2F pathway.

## Main

The E2F pathway is a transcriptional network that constitute a cardinal point of cell division and is essential to life. The E2F gene expression programs are highly conserved during evolution and act at the crossroads of cell proliferation, differentiation, apoptosis and stress responses to promote or halt the cell cycle. A family of eight E2F transcription factors work cooperatively or antagonistically to orchestrate the expression of genes necessary for DNA replication and cell cycle progression. Hence, an intricate balance between positive and negative regulators and feedback loops govern the E2F pathway and the cell proliferative capacity^1,2^. The E2F circuitry become perverted upon loss of tumor suppressors or activation/overexpression of oncogenes, both of which underlie tumor initiation and progression. On the other hand, FOXK1 and FOXK2 transcription factors, members of the Forkhead box (FOX) family, are known to regulate autophagy^6^, aerobic glycolysis^3^, insulin response^5^ and mTOR signaling^4^. However, evidence suggests that these factors might exert specific functions during cancer development and progression^14–20^. For instance, amplification of FOXK1 correlates with increased cell proliferation, as well as cancer progression^21^. In contrast, confounding results have been obtained on FOXK2 dysregulation in cancer^7^. As members of the FOX family, FOXK1 and FOXK2 contain a forkhead domain that mediates DNA binding^22^. These factors also contain a forkhead-associated (FHA) domain, exclusive to this family, which confers mutually exclusive interactions of FOXK1 or FOXK2 with phosphorylated BAP1^23^. A long-standing question regarding FOXK1 and FOXK2 is how these factors exert shared or distinct functions in coordinating biological processes. Here, we describe an important link between FOXK1/2 and the E2F pathway and reveal O-GlcNAcylation of FOXK1, but not FOXK2, as a molecular switch that distinctly promote cell proliferation and oncogenesis.

## FOXK1, but not FOXK2, promotes cell proliferation and is a potent oncogene

We first examined FOXK1 and FOXK2 mRNA levels in normal and cancer tissues, noting a general trend towards higher expression in tumors for both transcription factors (**Extended Data Fig. 1a and b**). Interestingly, the expression of FOXK1 closely correlated with that of FOXK2 in normal tissues when compared to other related FOX genes (**Extended Data Fig. 1c**). Moreover, the correlation between FOXK1 and FOXK2 becomes considerably weaker in cancer tissues. High levels of FOXK1 mRNA expression is associated with poor patient survival, while no association between FOXK2 expression levels and patient survival outcome was observed (**Extended Data Fig. 1d**).

To investigate the potential oncogenic properties of FOXK1/2, we first sought to explore the impact of their enforced expression in the context of normal human cell cycle progression. Notably, late passage IMR90 primary fibroblasts expressing FOXK1 become smaller and grow faster than empty vector or FOXK2 conditions (**Fig. 1a-c**). We then synchronized IMR90 cells expressing FOXK1 or FOXK2, with a combination of contact inhibition and serum deprivation to induce cell cycle exit, and followed cell cycle re-entry by re-plating the arrested cells at low density. FACS analysis showed that cells overexpressing FOXK1, but not FOXK2 or empty vector, can rapidly engage the S phase (**Fig. 1d**), consistent with an increased number of EdU positive S phase cells (**Fig. 1e**). Moreover, following three to four weeks of culture post-viral transduction, we observed a lower number of senescence-associated β-galactosidase (SA-β-gal)-positive cells in FOXK1 expression conditions comparatively to those of FOXK2 or empty vector (**Fig. 1f**), suggesting an extended replicative capacity of normal cells upon expression of FOXK1. Furthermore, immunostaining for PML bodies, known to be associated with cell senescence^24,25^, indicated that FOXK1-expressing cells present fewer number of senescence–associated PML bodies compared to FOXK2 or empty vector conditions (**Fig. 1g**). When we computed the numbers of IMR90 cells based on FOXK1 and FOXK2 expression levels (low versus high immunofluorescence signal intensity), we noticed that cells with higher FOXK1 signal intensity contains fewer numbers of PML foci per cell, consistent with our results that elevated FOXK1 expression delays the induction of cellular senescence (**Fig. 1h**). In contrast, the opposite results were observed for FOXK2, as higher number of PML bodies per cell correlates with higher FOXK2 expression levels (**Fig. 1h**). Of note, we also observed an increase in cellular proliferation following expression of FOXK1, but not FOXK2, in various cancer cells including osteosarcoma (U2OS) and colorectal carcinoma (HCT116) (**Extended Data Fig. 1e**).

**Figure 1:**
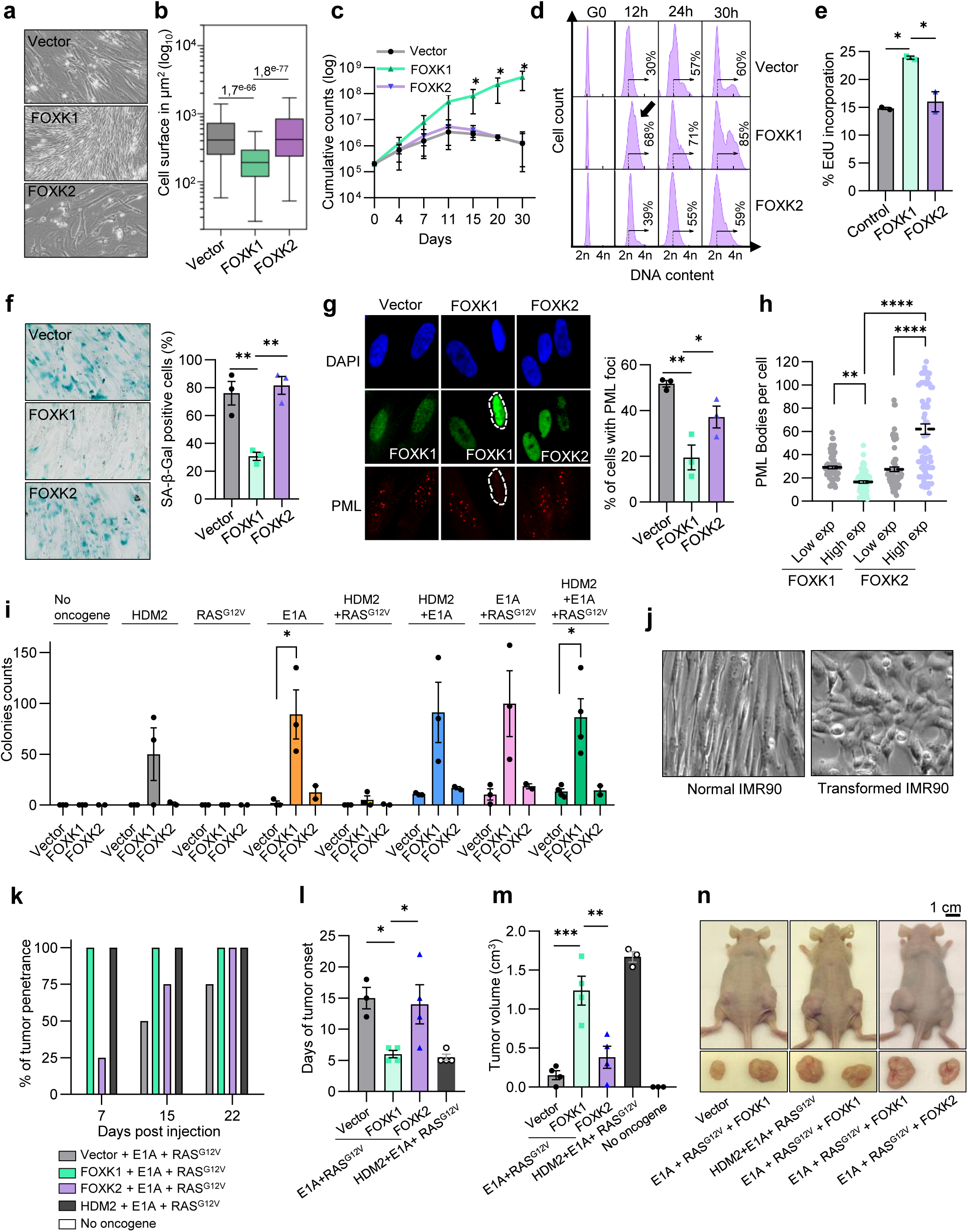
FOXK1 promotes cell proliferation and delays cellular senescence. **a, b**) Phase contrast imaging and cell size of IMR90 cells stably expressing empty vector, FOXK1 or FOXK2. The results are representative of more than 4 experiments. **c**) Cell counts of IMR90 cells expressing empty vector, FOXK1 or FOXK2. Data points are represented as a cumulative count (n=3). **d**) FACS analysis of cell cycle following synchronization and release of IMR90 cells expressing empty vector, FOXK1 or FOXK2. The percentage indicates the number of cells moving towards S/G2. The results are representative of three independent experiments. **e**) Analysis of EdU incorporation by immunofluorescence and cell counting of IMR90 cells expressing empty vector, FOXK1 or FOXK2 (n=2). **f**) Senescence-associated β-galactosidase staining of IMR90 cells expressing empty vector, FOXK1 or FOXK2. Cells stained in blue were counted and used to calculate the percentage of senescent cells (n=3). **g**) IMR90 cells expressing FOXK1 or FOXK2 were fixed for immunofluorescence staining of PML bodies (n=3). Control and FOXK1 expressing cells were stained with anti-FOXK1 antibody, FOXK2 expressing cells were stained with anti-FOXK2 antibody. Cells displaying PML bodies in each condition were counted and plotted in the right panel. **h**) Quantification of the number of PML bodies in cells with high or low expression of FOXK1 or FOXK2. **i**) Cell colony counting of IMR90 cells overexpressing empty vector, FOXK1 or FOXK2 along with different combinations of oncogenes. **j**) Representative images of normal versus transformed cells. **k**) Tumor penetrance of IMR90 cells expressing RAS^G12V^ and E1A and either empty vector, FOXK1 (n=5) or FOXK2 (n=2). The same number of cells were injected in the flank of nude mice. The experiment was terminated when the mice reach the limit point. **l**) Tumor latency of IMR90 cells expressing RAS^G12V^ and E1A, and either empty vector, FOXK1 or FOXK2 (n=3). **m**) Tumor volume of IMR90 cells expressing RAS^G12V^ and E1A and either empty vector, FOXK1 or FOXK2 at the end of the experiment (n=3). **n**) Images of the tumors before and after extraction for final size measurement. Statistics: Data are represented as mean ± SEM. \**P* < 0.05; \*\**P* < 0.01; \*\*\**P* < 0.001; \*\*\*\**P* < 0.0001. Student’s t-test (**b**, **c**, **e**, **i**). One-way ANOVA with Tukey’s multiple comparisons (**f**, **g**, **h**) or Dunnett’s (**l**, **m**).

To better define FOXK1 oncogenic proprieties, IMR90 cells were transformed by co-expressing different oncogenes in combination with FOXK1 or FOXK2. We also used RAS^G12V^, HDM2 and E1A, which is a classical combination of oncogenes known to transform normal human diploid fibroblasts when expressed together^26–28^. Expression of FOXK1 or FOXK2 along with RAS^G12V^ was not sufficient to induce colony formation (**Fig. 1i**). However, overexpressing FOXK1 with RAS^G12V^, HDM2 and E1A lead to a greater number of colonies compared to FOXK2 or empty vector conditions (**Fig. 1i**). These colonies acquire a rounded cell shape, in contrast to the fibroblast-like shape (**Fig. 1j)**, and can be expanded indefinitely. These results indicate that FOXK1 could further enhance the transformation potential of an otherwise potent oncogenic combination. In addition, expressing FOXK1 in cells with minimal combinations of oncogenes such as, HDM2 only, E1A only, HDM2 + E1A or E1A + RAS^G12V^, increased the number of colonies compared to controls (**Fig. 1i**). These cells have also acquired a tumorigenic potential and give rise to tumors when injected into immunodeficient mice. Notably, expressing FOXK1 with E1A and RAS^G12V^ combination could rapidly generate tumors with a high penetrance and a lower tumor latency than control cells or cells overexpressing FOXK2 (**Fig. 1k-n**). Moreover, tumors expressing FOXK1 are bigger than those expressing FOXK2 or control (**Fig. 1m,n**). Altogether, these results indicate that elevated expression of FOXK1 is observed in cancer and that this transcription factor constitutes a potent oncogene.

### FOXK1 is a major positive regulator of the E2F pathway

To define the mechanism underlying FOXK1-dependent oncogenic properties, transcriptomic (RNA-seq) analyses were conducted on IMR90 cells expressing either empty vector, FOXK1 or FOXK2. FOXK1 and FOXK2 transcripts were overexpressed approximately 4.7-fold and 14.5-fold respectively, compared to the empty vector condition (**Extended Data Fig. 2a**). Furthermore, we analyzed samples from TCGA cancer datasets, comparing those with the highest (top 10%) and lowest (bottom 10%) expression levels of FOXK1 or FOXK2. We observed that, on average, FOXK1 transcript counts varied about 8-fold between the two groups (high versus low), while FOXK2 transcript could varied around 7-fold (**Extended Data Fig. 2b**). These finding indicate that FOXK1 overexpression levels, observed in IMR90 cells, are reflective of the variations found in cancer contexts. Importantly, FOXK1 expression in IMR90 cells lead to differential expression of more than 2,000 genes with 902 upregulated and 1,112 downregulated compared to control, as well as about 475 genes with 286 upregulated and 189 downregulated when comparing FOXK1 to FOXK2 (**Fig. 2a and Extended Data Fig. 2c**). Gene ontology (GO) analysis revealed that FOXK1 is linked to several pathways regulating DNA replication and cell cycle progression (**Fig. 2b**). Notably, gene set enrichment analysis (GSEA) revealed that FOXK1 expression results in the activation of the E2F pathway (**Fig. 2b, c)**. Of the 200 E2F-regulated genes, we observed that 90 (45%) were increased in FOXK1-expressing cells, thus linking the enhanced cell proliferation to the upregulation of E2F target genes (**Fig. 2d**). In contrast, genes upregulated in FOXK2-overexpressing cells, compared to FOXK1, were associated with developmental processes and cellular differentiation (**Extended Data Fig. 2c**, cluster 3 and 4). Moreover, genes differentially regulated in FOXK2-overexpressing cells, compared to control condition, were associated with cell differentiation, migration, and adhesion (**Extended Data Fig. 2d,e**). FOXK1 and FOXK2 were previously shown to repress the autophagy pathway^6^ and, accordingly, we observed that genes associated with the regulation of autophagy were enriched in control cells compared to FOXK1 and FOXK2 (**Extended Data Fig. 2f**). Thus, FOXK1 and FOXK2 control overlapping and specific transcriptional programs in cells. In keeping with FOXK1 regulation of the E2F pathway, our transcriptomic analysis showed upregulation of E2F1 itself, as well as several E2Fs target genes such as FOXM1, cyclin A2, cyclin B1/2, CDC25C and MCM3 in FOXK1-overexpressing cells compared to FOXK2. We validated that FOXK1 promotes the expression of E2F1 and some of its known targets such as cyclin A2, MCM3 and CDC6 (**Fig. 2e**), which was also observed at the protein levels (**Fig. 2f**). Of note, FOXK2 overexpression results in a consistent induction of mRNA and protein levels of p21, a negative regulator of cell cycle (**Fig. 2e,f**).

**Figure 2:**
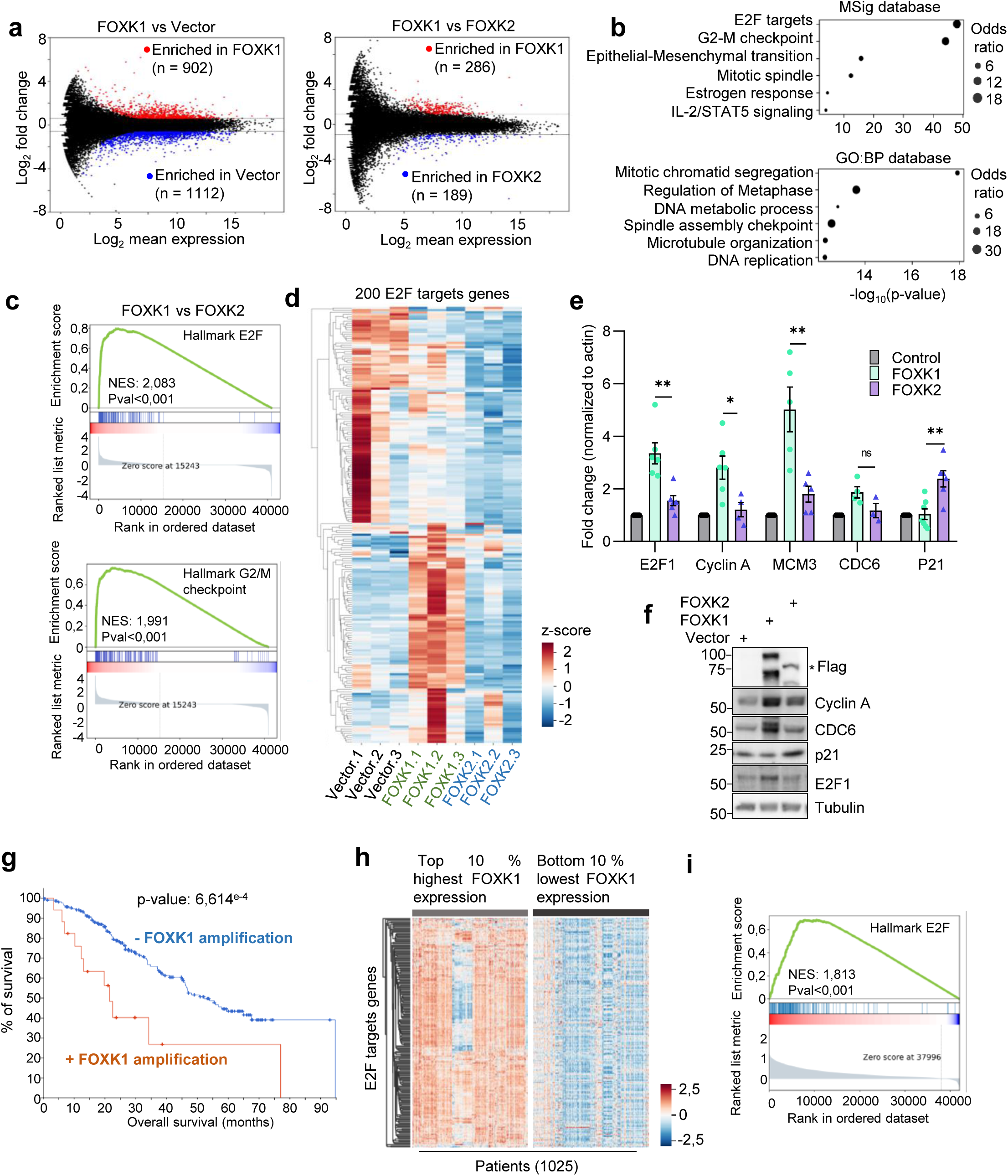
FOXK1 promotes the expression of E2F target genes. **a**) MA Plot representing the mean expression against the log fold change of genes when comparing FOXK1 with empty vector or FOXK1 with FOXK2 conditions. For each graph, genes in red are up regulated in FOXK1 condition. **b**) Gene ontology (GO) analysis using Enrichr (MSig and GO:BP databases) was performed on genes differentially regulated between FOXK1 and FOXK2 conditions. Odds ratio takes into account the number of input genes overlapping with the annotation set, the number of gene in the annotation set, the total number of genes in the input and the total number of genes in the human genome. See methods for details on computation. **c**) Gene set enrichment analysis (GSEA) performed on genes deregulated in FOXK1 compared to FOXK2 condition. **d**) Heatmap representing the transcript count of E2F target genes defined by the hallmark of molecular signatures database (200 genes) in control, FOXK1 and FOXK2 conditions. Transcript counts were normalized using z-score and presented as heatmap. **e**) Validation of RNA-seq data by quantifying mRNA of genes differentially regulated by qRT-PCR. Student’s t-test was performed. **f**) Western blotting showing increased expression of some E2F targets following FOXK1 or FOXK2 overexpression. **g**) Kaplan-Meier survival curve of TCGA cancer patients with or without FOXK1 amplification (cbioportal). **h**) Heat map of the 200 E2F target genes transcript counts (z-score) from TCGA cancer patients segregated between samples with the highest (top 10%) or lowest (bottom 10%) FOXK1 expression. **j**) GSEA analysis performed on genes differentially expressed when comparing TCGA samples with the highest versus the lowest expression of FOXK1.

FOXK1 amplification is observed in many solid cancers and its amplification is associated with decreased survival of patients (**Fig. 2g**). Next, we extracted mRNA expression from TCGA database, segregating it into two groups: the top 10% displaying the highest FOXK1 levels and the bottom 10% with the lowest FOXK1 expression. Samples with elevated FOXK1 mRNA showed pronounced expression of the 200 E2F target genes (**Fig. 2i**). Moreover, our differential gene expression analysis revealed a strong association between FOXK1 expression and the E2F pathway in cancer (**Fig. 2j**).

Our results indicate that FOXK1 expression is associated with the induction of the E2F pathway and could explain why cells are able to grow faster and are more susceptible to oncogenic transformation upon enforced expression of this transcription factor. This association is further mirrored in cancer tissues where samples with the highest levels of FOXK1 transcripts exhibit a pronounced activation of the E2F pathway, reinforcing the link between FOXK1 expression and oncogenesis.

### Pervasive occupancy of FOXK1 and FOXK2 of the E2F genomic circuit

To gain further insights into how FOXK1 and FOXK2 regulate gene expression, by notably discerning their common and specific target genes, we analyzed their genome occupancy in several cell lines using ChIP-seq. Remarkably, endogenous FOXK1 and FOXK2 exhibited similar chromatin recruitment patterns in IMR90 cells, co-localizing predominantly with the same promoters (**Fig. 3a,b**). ChIP-seq performed on IMR90 cells overexpressing Flag-tagged forms of FOXK1 and FOXK2 showed similar enrichment patterns as the corresponding endogenous proteins (**Extended Data Fig. 3a**). Next, gene ontology (GO) analysis of FOXK1/2 occupied promoters in these cells revealed several cellular processes linked to FOXK1/2 functions, including the E2F pathway (**Fig. 3c**). Further analysis of FOXK1 and FOXK2 genome occupancy across additional model cell lines indicated a consistent proportion of binding events in promoter regions with no redistribution of FOXK1 or FOXK2 binding sites upon their overexpression (**Fig. 3d**). Promoters commonly targeted by FOXK1 in U2OS, K562, and IMR90 (7,312 in total) were also associated with E2F and cell cycle regulation, hinting at a conserved role across different cell types (**Fig. 3e,f**, and **Extended Data Fig. 3b**). Importantly, genes upregulated by FOXK1 in IMR90 were associated with the binding of FOXK1 and FOXK2 on their promoters (273/289) (**Fig. 3g,h** and **Extended Data Fig. 3c**). We also analyzed distal regions (more than 1kb away, upstream and downstream from TSS) around these promoters and identified 1,193 regions occupied by FOXK1/2. Interestingly, FOXK1/2-bound promoters were enriched in E2F DNA binding motifs, while distal regions bound by FOXK1/2 were associated with other types of motifs such as Fra1/ATF3/AP-1 and CTCF, in addition to FOXK1 DNA binding motifs (**Fig. 3i**). Thus, FOXK1 and FOXK2 might link distal to promoter regions to orchestrate E2F gene expression programs. Importantly, while FOXK1 and FOXK2 are found on the same gene regulatory regions, only FOXK1 expression is associated with the induction of the E2F pathway suggesting differential regulation between these transcription factors at these genomic loci.

**Figure 3:**
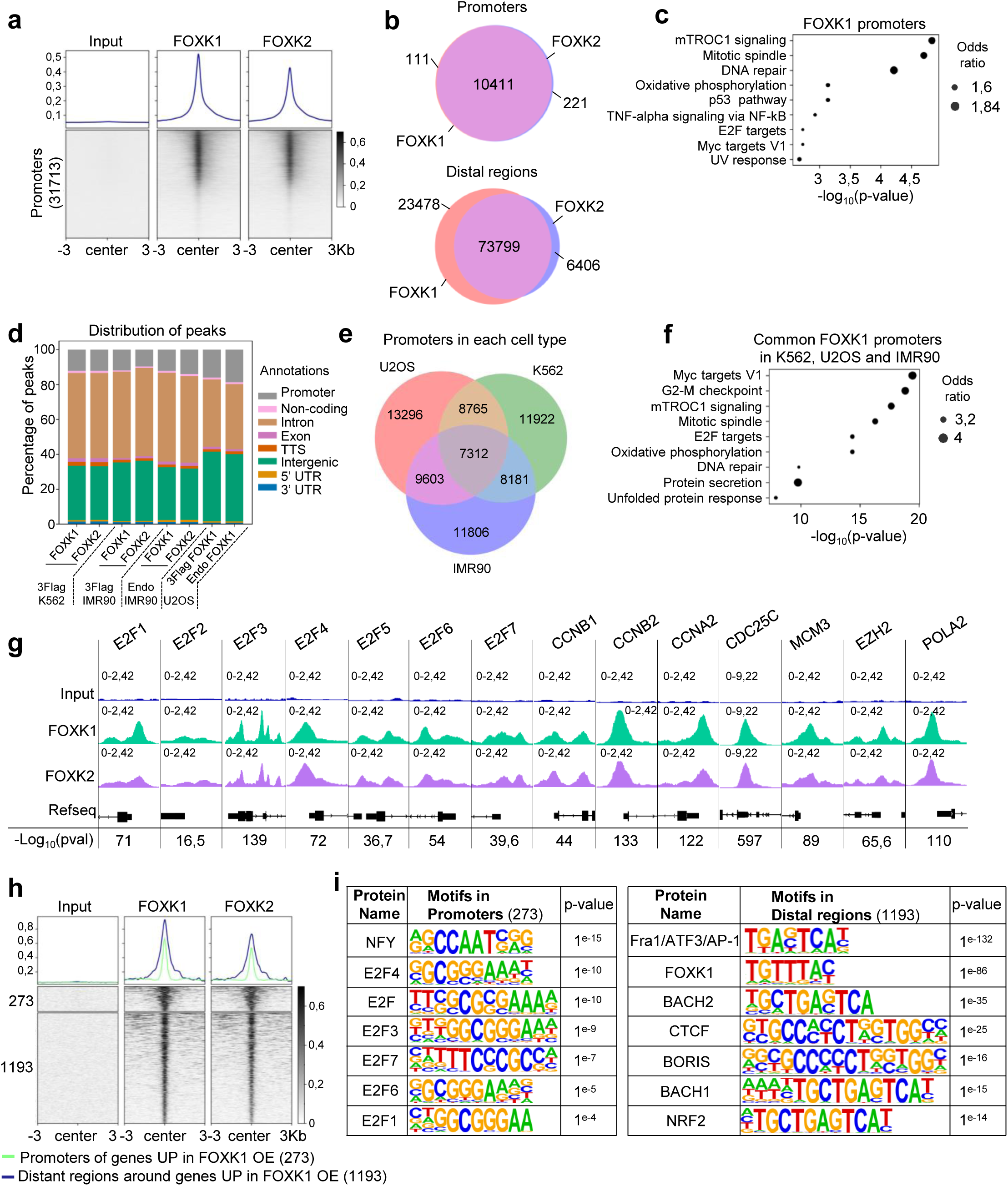
FOXK1 and FOXK2 occupy the same regulatory regions on chromatin. **a**) Heatmap and profile representing the occupancy of endogenous FOXK1 and FOXK2 on gene promoter regions. Promoter regions were obtained from HOMER (31713) and peaks were centered within 6kb (-/+ 3 kb) distance and oriented based on RefSeq direction. **b**) Venn diagram representing overlapping peaks in promoters and distal regions between endogenous FOXK1 and FOXK2 in IMR90 cells. The peaks were called with MACS2 with a p-value of 10^^-5^. **c**) Gene ontology (GO) analysis performed on promoters containing FOXK1. **d**) Bar-plot representing the repartition of endogenous (endo) and exogenous (3 Flag tagged) FOXK1 and FOXK2 ChIP-seq peaks on the genome of K562, IMR90 or U2OS cells. **e**) Venn diagram showing intersecting promoters containing FOXK1 (Flag ChIP-seq) in IMR90, K562 and U2OS cells. **f**) GO analysis performed on common 7312 promoters containing FOXK1 in IMR90, K562 and U2OS cells. **g**) Visualization of FOXK1 and FOXK2 occupancy on promoters of E2Fs and some of their target genes. Peaks p-value, called using MACS2, are shown under the gene body (Refseq) track. Peaks signal intensity is shown on the y axis. **h**) Occupancy of FOXK1 at promoters of 273 genes identified being differentially expressed in RNA-seq in IMR90 cells overexpressing (OE) FOXK1 compared to FOXK2. The 1193 distal regions were identified by considering peaks upstream or downstream promoters at a distance greater than 1kb away from TSS. **i**) Motif analysis was performed on promoters or distal regions indicated in panel h.

### FOXK1, but not FOXK2, is modified by O-GlcNAcylation

FOXK1 and FOXK2 are mutually exclusive partners of the BAP1 epigenetic complexes containing multiple co-factors and enzymes including the O-Linked β-N-Acetylglucosamine transferase (OGT), which mediates protein O-GlcNAcylation, a post-translational modification that regulate cellular metabolism and cell proliferation^29–35^. First, we tested whether FOXK1 or FOXK2 could be O-GlcNAcylated. Transient expression of FOXK1 or FOXK2 in the presence of OGT lead to the detection of a O-GlcNAcylation signal on immunoprecipitated FOXK1, but not FOXK2 (**Fig. 4a**). This signal is not observed following the expression of the catalytic dead (CD) form of OGT. Depletion of OGT expression using siRNA resulted in the ablation of the O-GlcNAcylation signal of endogenous FOXK1 (**Fig. 4b**). FOXK1 O-GlcNAc levels could be reliably increased by treatment with the OGA inhibitor PUGNAc, or decreased with the OGT inhibitor, OSMI-4 in IMR90 and other cell types (**Fig. 4c and Extended Data Fig. 4a**). Of note, modulation of cellular O-GlcNAc levels did not alter FOXK1 subcellular localization (**Extended Data Fig. 4b**). FOXK1 O-GlcNAcylation signal is directly linked to glucose availability, since it decreases under conditions of cell starvation (HBSS) or following incubation in glucose-free media, and increases upon gradual addition of glucose (**Fig. 4c,d**). Moreover, as FOXK1 regulates E2F expression, we sought to determine whether FOXK1 O-GlcNAcylation is modulated during the cell cycle. U2OS cells were synchronized by serum deprivation (**Extended Data Fig. 4c**), while primary human lung fibroblasts (HLF) and IMR90 were arrested through contact inhibition (**Fig. 4e and Extended Data Fig. 4d**). Endogenous FOXK1 was then immunoprecipitated at different times following release from cell cycle arrest. Interestingly, FOXK1 O-GlcNAcylation and CDC6 expression reached their maximum levels at the same time point, suggesting that activation of E2F-dependent transcription is concomitant with FOXK1 O-GlcNAcylation (**Fig. 4e and Extended Data Fig. 4c,d**), and highlighting a potential role of O-GlcNAcylation in regulating FOXK1 activity during the cell cycle. Of note, FOXK1 O-GlcNAcylation is decreased during differentiation of 3T3L1 adipocytes, supporting the notion that this post-translational modification is associated with FOXK1-dependent stimulation of cell proliferation (**Extended Data Fig. 4e**).

**Figure 4:**
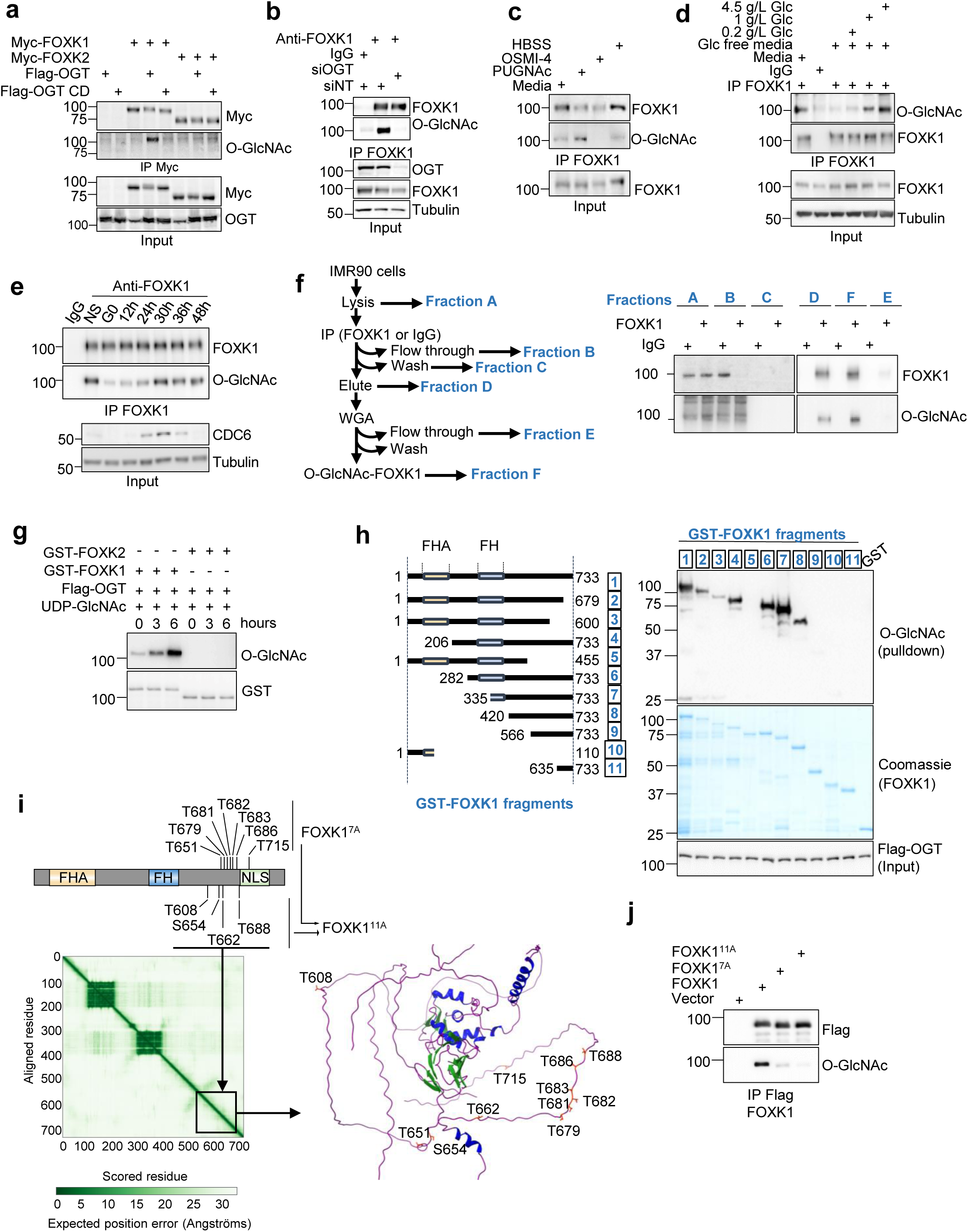
FOXK1, but not FOXK2, is modified by O-GlcNAcylation. **a**) HEK293T cells were transfected with constructs expressing Myc-FOXK1 or Myc-FOXK2 in the presence of OGT WT or OGT catalytically dead (CD) mutant. Myc immunoprecipitation was performed, and levels of O-GlcNAcylation were detected using an anti-O-GlcNAc specific antibody (n=2). **b**) Immunoprecipitation of endogenous FOXK1 was performed on U2OS cell extracts transfected with siRNA targeting OGT (siOGT) or non-target siRNA as a control (siNT) (n=3). **c**) Immunoprecipitation of endogenous FOXK1 and analysis of O-GlcNAcylation in IMR90 cells treated with either; modified Hanks’ Balanced Salt Solution (HBSS) (no glucose or amino acids), OGA inhibitor (PUGNAc) or OGT inhibitors (OSMI-4) (n=3). **d**) Immunoprecipitation and analysis of endogenous FOXK1 O-GlcNAcylation in IMR90 cells treated with glucose free media or gradually supplemented with increasing concentrations of glucose (n=3). **e**) Immunoprecipitation and analysis of endogenous FOXK1 O-GlcNAcylation in IMR90 cells synchronized by contact inhibition and released at low density in fresh medium (n=3). **f**) Immuno-depletion and analysis of endogenous FOXK1 O-GlcNAcylation in IMR90 cells. Cellular extracts from IMR90 were used for FOXK1 immunoprecipitation. Eluted proteins were then incubated with WGA coated beads and FOXK1 O-GlcNAcylation levels were analyzed by western-blotting (n=3). **g**) In vitro O-GlcNAcylation was performed on recombinant GST-FOXK1 or GST-FOXK2 with recombinant His-OGT-Flag. The reaction was stopped at different time points to detect protein O-GlcNAcylation levels (n=3). **h**) Left: recombinant FOXK1 fragments are schematically represented and numbered. Right: in vitro O-GlcNAcylation was performed on recombinant FOXK1 fragments to map the region containing residues modified by O-GlcNAc (n=3). **I**) Top; schematic representing the identification of O-GlcNAc sites on FOXK1 as determined by mass spectrometry (MS) analysis. Mutant FOXK1^7A^ contains seven threonine mutated to alanine, whereas mutant FOXK1^11A^ contains all the eleven sites mutated to alanine. Bottom; FOXK1 structure predicted by Alphafold. The region highlighted is expected to be unstructured. Right; Visual representation of this region with the position of residues targeted by O-GlcNAcylation are shown on the predicted protein structure. **j**) Immunoprecipitation of Flag-tagged versions of FOXK1, FOXK1^7A^, or FOXK1^11A^ from stable IMR90 cell extracts and detection of O-GlcNAcylation levels (n=3).

Our results indicate that FOXK1 O-GlcNAcylation is dependent on the metabolic state of the cell as well as it cell cycle state. Thus, we reasoned that FOXK1 molecules exist under O-GlcNAcylated or non-O-GlcNAcylated forms. Alternatively, FOXK1 molecules might be O-GlcNAcylated on multiple sites, but with different degrees of modifications. To further determine the extent of FOXK1 O-GlcNAcylation in exponentially proliferating cells, we first immunodepleted endogenous FOXK1 from IMR90 cell extracts and subsequently incubated the immunopurified FOXK1 on wheat germ agglutinin (WGA) coated beads to capture the fraction of O-GlcNAcylated FOXK1. Fractions were collected at all steps including the flow through and probed for O-GlcNAc and FOXK1 (**Fig. 4f**). Interestingly, we observed that nearly all endogenous FOXK1 is O-GlcNAcylated. Thus, FOXK1 is likely to contain multiple sites whose extent of O-GlcNAcylation varies depending on cellular states.

To identify FOXK1 O-GlcNAcylation region, *in vitro* O-GlcNAcylation assays were performed using recombinant GST-FOXK1 or GST-FOXK2, OGT and UDP-GlcNAc. First, we confirmed that FOXK1, but not FOXK2, is O-GlcNAcylated by OGT *in vitro* (**Fig. 4g**). In addition, *in vitro* O-GlcNAcylation on recombinant fragments of FOXK1 showed that O-GlcNAcylation occurs in the C-terminal region of the protein, with FOXK1 fragment 1 to 455 amino acids losing its ability to be modified by OGT (**Fig. 4h**). Interestingly, while the O-GlcNAcylation of FOXK1 occurs at the C-terminus, the OGT-FOXK1 interaction also involves the N-terminal part of the protein (**Fig. 4h** and **Extended Data Fig. 5a**). Next, we sought to identify the FOXK1 amino acid residues modified by O-GlcNAcylation. FOXK1 was overexpressed along with OGT in HEK293T and a large-scale immunopurification was performed to ensure high protein recovery for mass spectrometry. We identified seven residues in the C-terminal region that are modified by O-GlcNAcylation (**Extended Data Fig. 5b**). We generated an expression construct, FOXK1^7A^, by mutating the seven residues targeted by O-GlcNAcylation to alanine (**Fig. 4i**). We expressed the FOXK1^7A^ mutant in IMR90 and other cell types and noticed that this mutant has reduced O-GlcNAcylation signal, but a residual modification signal is still observed (**Fig. 4j and Extended data Fig. 5a-c**). We then purified FOXK1^7A^ and identified four additional residues that are modified by O-GlcNAc. All sites were found in an unstructured region of the C-terminal of the protein (**Fig. 4i**). We therefore produced a second mutant, FOXK1^11A^, where we mutated the remaining four serine or threonine to alanine (**Fig. 4i**).

**Figure 5:**
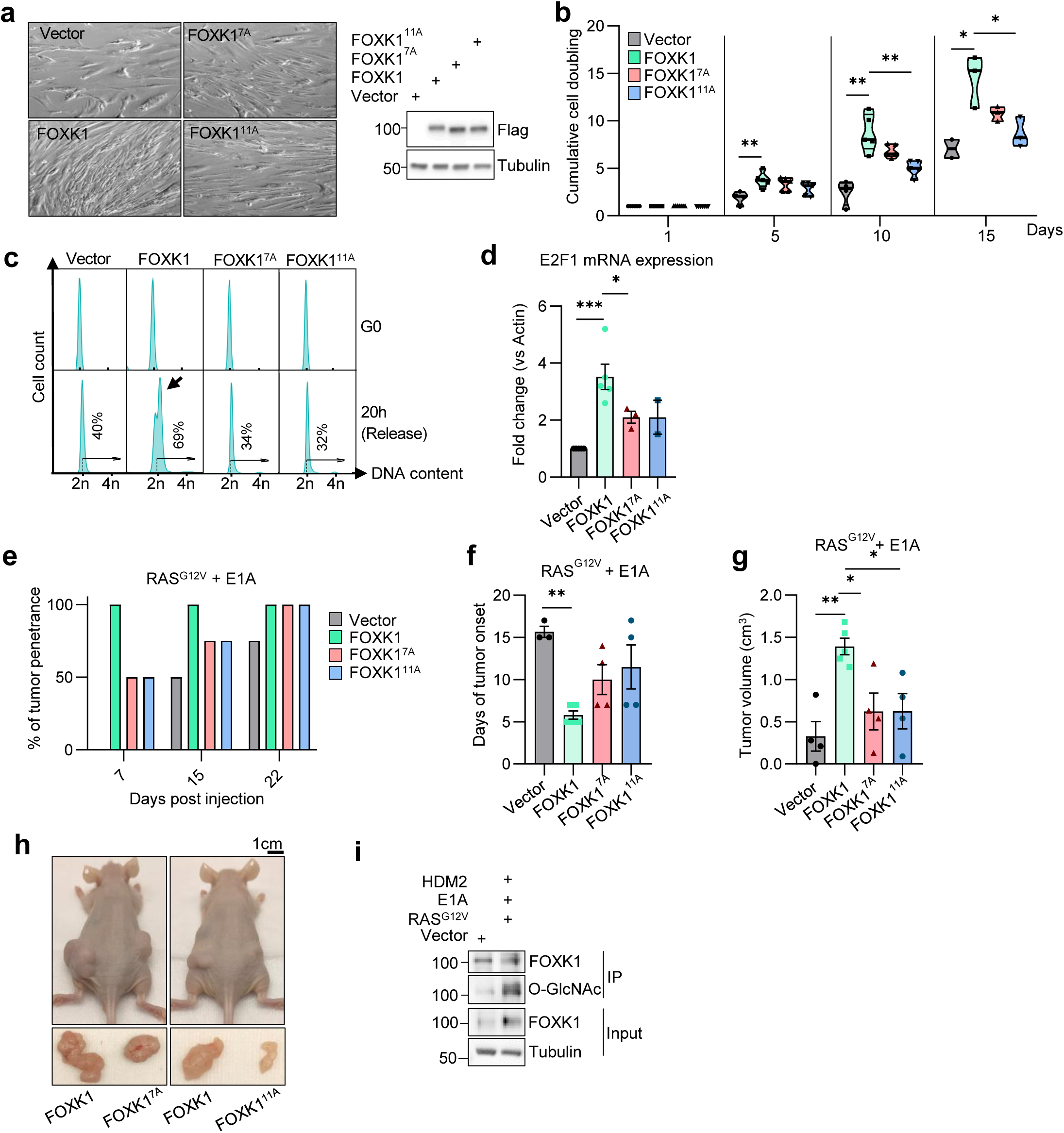
O-GlcNAcylation regulates FOXK1 oncogenic proprieties. **a**) Phase contrast and immunoblotting of IMR90 cells overexpressing FOXK1, FOXK1^7A^ or FOXK1^11A^. **b**) IMR90 cells overexpressing FOXK1, FOXK1^7A^ or FOXK1^11A^ were counted over 15 days. Data are represented as a cumulative cell doubling plot (n=3). **c**) IMR90 cells stably expressing FOXK1, FOXK1^7A^, FOXK1^11A^ or empty vector, were blocked in G0 by contact inhibition, and released by plating at low density in fresh medium to monitor cell cycle progression by FACS analysis. Results are representative of three independent experiments. **d**) E2F1 mRNA quantification by RT-qPCR in IMR90 cells overexpressing FOXK1, FOXK1^7A^ or FOXK1^11A^. **e**) Tumor penetrance of xenograft performed with IMR90 cells expressing RAS^G12V^ with E1A in combination with either empty vector, FOXK1, FOXK1^7A^ or FOXK1^11A^ (n=4). **f**) Tumor latency representing the time between cell engraftment and appearance of tumors that reached at least 0.1 cm^3^. **g**) Tumor volume was calculated at the end of the experiment. All tumors were harvested at the same time when the fastest growing tumors reached 1.7 cm^3^ (n=4). **h**) Representative images of tumors before and after extraction. **i**) Immunoprecipitation of endogenous FOXK1 from normal or transformed IMR90 (combination of RAS^G12V^ with E1A and HDM2) to evaluate FOXK1 O-GlcNAcylation levels. Representative of three independent experiments. Data are represented as mean ± SEM (**d, f and g**). Multiple t-test (**b**). One-way ANOVA with Dunnett’s multiple comparisons **(d**, **f**, **g**). \**P* < 0.05; \*\**P* < 0.01; \*\*\**P* < 0.001.

FOXK1^11A^ showed a more pronounced decrease of O-GlcNAcylation comparatively to the FOXK1^7A^ mutant in IMR90 and other cell lines (**Fig. 4j and Extended data Fig. 6a-c**). Of note, we observed only marginal changes of FOXK1 O-GlcNAcylation levels when mutating individual residues (**Extended Data Fig. 6d**). FOXK1 O-GlcNAcylation levels decrease only upon mutation of multiple sites, and the strongest decrease observed when mutating all eleven residues identified (**Extended Data Fig. 5a-c and Extended data. Fig. 6e**). Loss of FOXK1 O-GlcNAcylation had no effect on protein stability and protein subcellular localization (**Extended Data Fig. 6f,g**). Taken together, O-GlcNAcylation specifically targets FOXK1, but not FOXK2, which could constitute a molecular switch underlying their differential functions.

**Figure 6:**
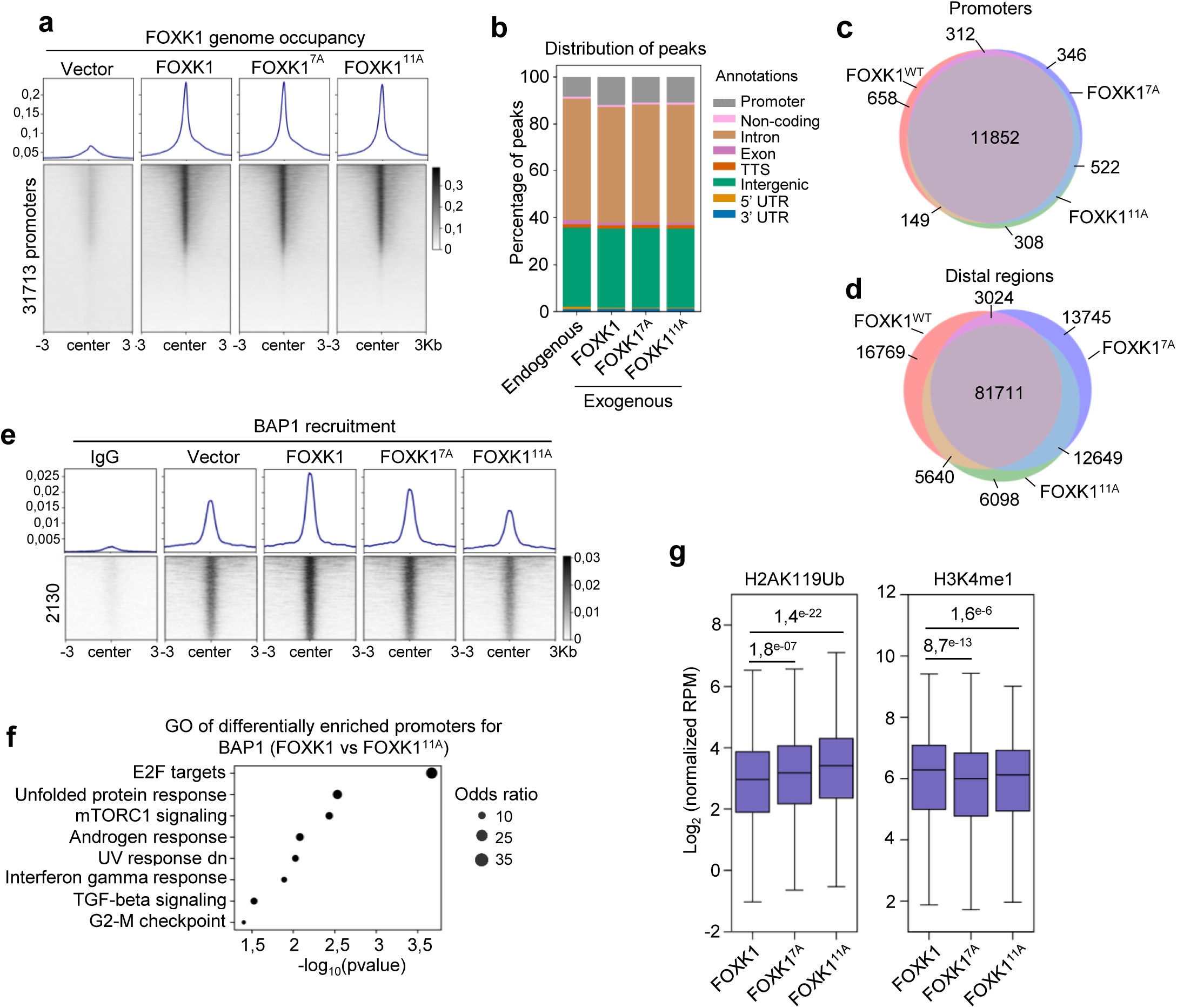
FOXK1 O-GlcNAcylation regulates its transcriptional function on chromatin. **a**) Chromatin occupancy of Flag-tagged FOXK1, FOXK1^7A^ and FOXK1^11A^ on all human promoters in IMR90 cells. **b**) Bar-plot representing the repartition of endogenous FOXK1 and exogenous (3 Flag tagged) FOXK1, FOXK1^7A^ and FOXK1^11A^ on the genome of IMR90 cells assessed by ChIP-seq. **c-d**) Venn diagram depicting the overlap in chromatin occupancy between FOXK1, FOXK1^7A^ and FOXK1^11A^ at promoters and at distal regions in IMR90 cells. **e**) Differential recruitment of BAP1 in IMR90 cells overexpressing FOXK1, FOXK1^7A^ or FOXK1^11A^. Regions were identified by comparing BAP1 recruitment in FOXK1 with BAP1 recruitment in FOXK1^11A^. Technical replicate were merged for visualization. **f**) GO analysis performed on promoters (249) differentially enriched for BAP1 between FOXK1 and FOXK1^11A^. **g**) Boxplot representing H3K4me1 and H2AK119ub normalized reads per million (RPM) in IMR90 cells expressing FOXK1, FOXK1^7A^ or FOXK1^11A^ on regions with differential BAP1 recruitment.

### FOXK1 O-GlcNAcylation is required for cell proliferation and tumor progression

We sought to determine the potential contribution of O-GlcNAcylation to FOXK1 oncogenic properties. Expression of FOXK1^7A^ and FOXK1^11A^ in IMR90 cells led to reduced cell proliferation comparatively to FOXK1 (**Fig. 5a,b**). In addition, synchronized cells expressing FOXK1^7A^ and FOXK1^11A^ progressed more slowly into S phase compared to FOXK1 (**Fig. 5c**). FOXK1 mutants with impaired O-GlcNAcylation failed to induce E2F1 expression as efficiently as the wild type form, indicating that O-GlcNAcylation modulates the ability of FOXK1 to stimulate E2F genes (**Fig. 5d**). To further determine whether the loss of O-GlcNAcylation can also impact FOXK1 oncogenic properties, we performed oncogenic transformation using IMR90 cells expressing RAS^G12V^ with E1A in combination with either FOXK1, FOXK1^7A^ or FOXK1^11A^ (**Extended Data Fig. 7a**). A delayed onset of tumors with a longer latency period were observed in mice engrafted with cells expressing FOXK1^7A^ or FOXK1^11A^ compared to those engrafted with cells expressing FOXK1 (**Fig. 5e,f**). Cells overexpressing FOXK1 developed tumors that reached the limit point faster than cells overexpressing FOXK1^7A^, FOXK1^11A^ (**Fig. 5g,h**). The oncogenic effect of FOXK1 O-GlcNAcylation was also observed with the minimal transforming combinations of HDM2, RAS^G12V^ or E1A with FOXK1 which could also lead to tumor formation in mice, although at a much lower penetrance than the HDM2+ E1A +RAS^G12V^ combination (**Fig. 1g and Extended data. 7b,c**).

Previous studies reported that elevated protein O-GlcNAcylation in cancer can sustain tumor cell proliferation and progression^36,37^. This raises the possibility that increased O-GlcNAcylation can further sustain the oncogenic effect associated with high FOXK1 expression. Consistent with this, transformed IMR90 cells (E1A + RAS^G12V^ + HDM2) displayed increased levels of FOXK1 O-GlcNAcylation compared to corresponding normal cells (**Fig. 5i**). To further determine the impact of FOXK1 O-GlcNAcylation on cancer cell proliferation and tumor progression, we first depleted endogenous FOXK1 in U2OS osteosarcoma cells using siRNA, which resulted in decreased cell proliferation (**Extended Data Fig. 7d**). This effect was rescued by expression of FOXK1 but not FOXK1^7A^ siRNA resistant constructs, and is associated with decreased mRNA levels of the E2F target gene, cyclin A (**Extended Data Fig. 7d,e**). To further assess if loss of FOXK1 O-GlcNAcylation could impact on tumor progression *in vivo*, we performed xenograft experiments using the prostate cancer cell line PC3, previously shown to exhibit high O-GlcNAc levels^38^. We first confirmed that overexpressing FOXK1^7A^, or FOXK1^11A^ in PC3 reduced cell proliferation compared to FOXK1 (**Extended Data Fig. 7f**). We then engrafted FOXK1-expressing cells in the flank of nude mice and observed reduced tumor growth with FOXK1 mutants impaired for O-GlcNAcylation (**Extended Data Fig. 7g,h**). In addition, expression of the FOXK1^7A^ and FOXK1^11A^ O-GlcNAcylation-defective mutants resulted in decreased E2F1 protein levels compared to FOXK1 (**Extended Data Fig. 7i**). Altogether, these findings illustrate that O-GlcNAcylation supports FOXK1 pro-oncogenic functions in promoting cellular transformation and tumor growth.

### FOXK1 O-GlcNAcylation promotes BAP1 recruitment to E2F target gene promoters

To investigate whether O-GlcNAcylation regulates FOXK1 recruitment to chromatin, we performed ChIP-seq in IMR90 cells, which revealed conserved peaks near promoters and distal regions among FOXK1, FOXK1^7A^ and FOXK1^11A^ (**Fig. 6a-d, Extended data Fig. 8a-b**). Additionally, ATAC-seq confirmed FOXK1 association with open chromatin, which remained unaffected following expression of its O-GlcNAcylation defective mutant (**Extended Data Fig. 8c**). FOXK1/2 were previously shown to recruit the histone H2AK119ub deubiquitinase BAP1 to chromatin and mediate transcriptional activities^32^. By comparing BAP1 recruitment in IMR90 cells expressing FOXK1 with those expressing FOXK1^11A^, we identified 2,130 regions showing reduced recruitment of BAP1 in FOXK1^11A^ (**Fig. 6e**). GO analysis on promoters (249) contained in these differentially enriched regions (2,130) revealed a strong association with E2Fs target genes (**Fig. 6f**). BAP1 was previously found to interact with and recruit the methyl-transferase MLL3, which is responsible for the deposition H3K4me1 at enhancers^40, 41^. Interestingly, IMR90 cells expressing FOXK1^7A^ or FOXK1^11A^ exhibited reduced levels of H3K4me1 on regions with reduced BAP1 recruitment (**Fig. 6g**). Conversely, H2AK119ub levels were increased in FOXK1^7A^ or FOXK1^11A^, correlating with the reduction of BAP1 recruitment in these conditions (**Fig. 6g**). This result suggest that loss of O-GlcNAcylation on FOXK1 perturbate the optimal chromatin configuration and hint to a potential mechanism for how O-GlcNAcylation of FOXK1 regulates transcription. Next, we questioned how BAP1 recruitment was affected on promoters of E2F target genes whose expression was induced by FOXK1 (**Fig. 3h**). Notably, despite the constant occupancy of FOXK1 regardless of its O-GlcNAcylation status, we observed decreased association of BAP1 with these regions for FOXK1^7A^ and FOXK1^11A^ comparatively to FOXK1 (**Extended Data Fig. 8d** and **Extended Data Fig. 9a**). Further, BAP1 was found to co-localize with BRD4 and H3K27Ac in IMR90^39^, as well as with H3K4me1 (**Extended Data Fig. 9a,b**), suggesting that these distal regions are enhancers. Additionally, we observed reduced levels of H3K4me1 and increased levels of H2AK119ub at both promoters and enhancers (**Extended Data Fig. 9b**). This findings imply that these regulatory regions are sensitive to FOXK1 O-GlcNAcylation, and that optimal transcriptional activity require the deposition of O-GlcNAcylation on FOXK1. Taken together, our data indicate that O-GlcNAcylation of FOXK1 regulates the optimal recruitment of BAP1 to chromatin. Abolishing O-GlcNAcylation leads to decreased BAP1 recruitment at promoters and surrounding enhancers of E2F target genes. This event is associated with a corresponding decrease of H3K4me1 and increase H2AK119ub, explaining the switch from transcriptional active to inactive chromatin states. Thus, O-GlcNacylated FOXK1 associates with BAP1 and promote E2F pathway and oncogenesis.

## Discussion

Our findings indicate that FOXK1 is a potent oncogene and a major regulator of the E2F pathway. We also revealed that FOXK1 oncogenic properties require O-GlcNAcylation, which could be an important general mechanism of tumorigenesis in human malignancies. Cancer cells are known to have increased activity of the glycolytic pathway which is thought to be a quick way to provide energy and building blocks required during fast cellular growth^42^. Perturbation of O-GlcNAcylation levels are also observed in cancer and different mechanisms were proposed to explain how increased protein O-GlcNAcylation is favorable for cancer development^43^. Previous studies demonstrated that FOXK1 regulates the glycolytic pathway and its overexpression promote glucose consumption and reprograming of cell metabolism to favor glycolysis^3,5^. FOXK1 could therefore increase glucose uptake to fuel the hexosamine biosynthetic pathway and promote the synthesis of UDP-GlcNAc and, as a result, further enhance FOXK1 O-GlcNAcylation. Because O-GlcNAcylation is dependent on glucose availability in cells, O-GlcNAcylation might be a mechanism to regulate FOXK1 activity depending on the state of cellular metabolism and cell microenvironment. We propose that, depending on glucose availability and cellular metabolism, the extent of FOXK1 O-GlcNAcylation on its multiple sites might serve as a rheostat to regulate its transcriptional activity on the E2F pathway and orchestrate cell proliferation.

We also found that FOXK1 and FOXK2 are recruited to the same genomic loci, but only FOXK1 was able to induce genes associated with the positive regulation of cell cycle. We showed that O-GlcNAcylation acts as a mechanism to specifically regulate FOXK1 transcriptional activities by targeting BAP1 recruitment to chromatin. Loss of FOXK1 O-GlcNAcylation reduced BAP1 recruitment to promoters and enhancers and is associated with an increased level of H2AK119ub, a mark associated with the negative regulation of transcription. In addition, H3K4me1 was decreased at a subset of promoters and enhancers associated with E2F targets. Thus, we propose that O-GlcNAcylation fine tune FOXK1 transcriptional activities by allowing optimal recruitment of BAP1. Paradoxically, although BAP1 is a tumor suppressor, it appears counterintuitive that the oncogenic functions of FOXK1 rely on BAP1. However, previous studies have indicated that BAP1 can promote the expression of E2F targets, and moreover, the oncogenic properties of KLF Transcription Factor 5 (KLF5) also appear to be mediated through BAP1 function^44^. It remains to be determined how the interplay between FOXK1 O-GlcNAcylation and BAP1 could orchestrate the E2F pathway to impact transcription and tumorigenesis.

## Methods

### Molecular DNA cloning and mutagenesis

Plasmids for expression of human Myc-OGT and Myc-OGT D925A catalytic inactive mutant (Myc-OGT CD) were previously described^45^. His-OGT-Flag was generated by subcloning the OGT cDNA into pET30a+ vector (Novagen^®^). siRNA-resistant human FOXK1 and FOXK2 cDNAs were synthesized into a pBluescript plasmid (Biobasic^®^) and subcloned into pENTR (Life technologies^®^). GST-, FLAG- and MYC-tagged, retroviral pMSCV-Flag/HA (Addgene, #41033) and pMSCV-3Flag (generated for this study) constructs were generated using the Gateway recombination system (Life technologies). GST-FOXK1 fragments were generated by PCR and subcloned into pENTR. FOXK1 single and multiple O-GlcNAc sites mutants including FOXK1^7A^ and FOXK1^11A^ mutants were generated with site directed mutagenesis or gene synthesis and subcloned in appropriate bacteria or mammalian expression vectors. We use pCMV-VSVG (#8454, Addgene) and HELPER (a gift from Dr. F.A. Mallette) to generate retroviral particles. The following retroviral vectors were used for cellular transformation: pWZL-hygro E1A (#18748, Addgene), pWZL-Blast RAS^GV12^ (#12277, Addgene), and pWZL-neo HDM2^46^ (a gift from Dr. F.A. Mallette). All DNA constructs were sequenced

### Alphafold structure prediction

The AlphaFold Protein Structure Database was used to retrieve human FOXK1 structural model (Uniprot: P85037-F1)^47,48^. Visualization of the structural model and the corresponding amino acids was generated using ChimeraX^49^. The side chains of the corresponding amino acid are shown as sticks. The O-GlcNAcylation sites are located within a C-terminal region with a per-residue model confidence score (pLDDT) below 50, likely corresponding to an unstructured region.

### Cell culture and treatments

Human lung fibroblasts (IMR90, CL-173), transformed human embryonic kidney cells (HEK293T, CRL32-16), transformed human embryonic kidney cells (Phoenix-AMPHO, CRL-3213), human osteosarcoma cells (U2OS, HTB96), chronic myelogenous leukemia (K562, CCL-243), prostatic adenocarcinoma (PC-3, CRL-1435), murine preadipocyte (3T3-L1, CL-173), human colon cancer (HCT116, CCL-247) and cervical carcinoma cells (HeLa, CCL-2) were purchased from ATCC^®^. Primary lung fibroblasts (HLFs) were obtained from Dr. Elliot Drobetsky (Montreal University). Cells were cultured at 37°C and 5% CO2, in Dulbecco’s modified Eagle’s medium (DMEM, Wisent^®^, 319-005-CL) supplemented with 10% foetal bovine serum (FBS, Wisent, 098150) or in 5% new-born calf serum (NBS, Sigma^®^, N4637) with 2% FBS. K562 cells were cultured in RPMI-1640 medium supplemented with 5% NBS. Media were supplemented with 4 mM L-Glutamine (Bioshop^®^, GLU02.500), 100 U/ml Penicillin (Biobasic, PB-0135) and 100 µg/ml Streptomycin (Bioshop, STP101.100). Cells were regularly tested for mycoplasma contamination by PCR and DAPI staining. For modulation of FOXK1 O-GlcNAcylation levels, cells were treated, in the complete culture medium, with 10 µM of the OGT inhibitor OSMI4, or 50 µM the OGA inhibitor PUGNAc or Thiamet G and harvested at the indicated times for immunoprecipitation or immunofluorescence. Cells were also incubated in a modified Hank’s Balanced Salt Solution (HBSS) containing no glucose or amino acids. For experiments with increasing glucose concentration, cells were incubated in glucose-free culture medium completed with 0 g/L, 0.2 g/L, 1 g/L or 4.5 g/L of glucose for the indicated times and harvested for immunoprecipitation and immunoblotting. For analysis of FOXK1 stability, cells were treated with 20 µM of MG132 (Sigma, C221) or 20 µg/ml of cycloheximide (Bioshop, CYC003.1) and harvested at the indicated times for immunoprecipitation or immunoblotting.

### Cell cycle synchronisation and flow cytometry Analysis

U2OS cells were grown to confluence and then serum starved for 24 hours. Cells were subsequently incubated in fresh media containing 20 % FBS for the indicated times before being harvested for immunoprecipitation and flow cytometry analysis. Primary human fibroblasts were grown to confluence and further cultured for 3 days. The cells were then serum starved for two days, trypsinized and replated in fresh medium before being harvested at the indicated times for immunoprecipitation and flow cytometry analysis. For flow cytometry analysis, the cells were harvested by trypsinization and fixed with 75 % ethanol. Following centrifugation, cells were incubated for 30 min at 37 °C in PBS containing 100 µg/ml RNase A (Biobasic, RB0473) for 30 min before DNA staining with 50 µg/ml propidium iodide. Cell DNA content was determined with a FACSCalibur™ flow cytometer (BD Biosciences^®^) and analyzed with the CellQuest™ Pro software (BD Biosciences).

### Cell differentiation

3T3L1 differentiation was done essentially as described before^31^. Exponentially proliferating cells were grown to confluence and then left at confluence for 48 hours before incubation in differentiation media (DMEM supplemented with 10% FBS, 4 mM L-Glutamine, 100 U/ml penicillin/streptomycin, 1 μM dexamethasone (Sigma, D-2915), 1 μg/ml insulin (Sigma, I5500) and 500 μM isobutylmethylxanthine (Sigma, I5879). Two days post-induction, the differentiation medium was changed for complete DMEM medium supplemented with 1 μg/ml insulin. Media were changed every 48 hours and cells were harvested at the indicated time points for immunoprecipitation or immunoblotting.

### Colony forming assay (CFA)

Similar numbers of U2OS, HCT116 or PC3 cells stably expressing the different constructs of FOXK1 or FOXK2 were seeded on 6 cm or 10 cm plates. Following 3 to 10 days of culture, the surviving colonies were washed with PBS and fixed with 3% paraformaldehyde (PFA) for 20 min. Cells were then washed with PBS once and stained with 0.2% crystal violet for 10 min. Following several washes with water, the plates were imaged and colonies counted.

### SA-β-gal activity assay

SA-β-gal activity assay was performed as previously described^50,51^. Briefly, cells were fixed with 0.5% glutaraldehyde (Sigma, G5882) in PBS for 15 min, then washed and kept in PBS (pH 6.0) containing 1 mM of MgCl_2_, for at least 24 hours. SA-β-gal staining was performed at 37°C using a solution containing X-Gal, potassium ferricyanide, potassium ferricyanide and MgCl_2_ in PBS (pH 6.0). Images were taken with an inverted microscope and the percentage of SA-β-gal positive cells was quantified in each condition.

### Antibodies

A rabbit polyclonal anti-FOXK2 antibody was generated and validated by RNAi. Mouse monoclonal anti-FOXK1 (G4, sc-373810), mouse monoclonal anti-BAP1 (C4, sc-28383), rabbit polyclonal anti-OGT (H300, sc-32921), mouse monoclonal anti-tubulin (B-5-1-2, sc-SC-23948), mouse monoclonal anti-CDC6 (180.2 sc-9964), rabbit polyclonal anti-FOXK1 (H140, sc-134550), mouse monoclonal anti-E2F1 (KH95, sc251), mouse monoclonal anti-cyclin A2 (6E6, sc-56299), mouse monoclonal anti-HSP90⍺/β (F8, sc-13119), mouse monoclonal anti-PML (PG-M3, sc-966), mouse monoclonal anti-HRAS (C-20, sc-520) were from Santa Cruz Biotechnologies^®^. Rabbit polyclonal anti-HCF-1 (A301-400A) was from Bethyl Laboratories^®^. Mouse monoclonal anti-Flag (M2), mouse monoclonal anti-Actin (MAB1501, clone C4) and rabbit polyclonal anti-GST (G7781) were from Sigma-Aldrich. Rabbit polyclonal anti-FOXK1 (MNF, ab-18196), monoclonal anti-O-Linked N-acetylglucosamine (RL2, ab2739), rabbit polyclonal anti-H3 (ab1791) were from Abcam^®^. Mouse monoclonal anti-Rb (4H1, 9309S), rabbit polyclonal anti-phospho Rb (S807/811, 9308), rabbit mono anti-USP10 (D7A5, 8501), rabbit polyclonal anti-Perilipin (D1D8, 9349) were from Cell Signaling^®^. Rabbit polyclonal anti-FABP4 (10004944) was from Cayman Chemical^®^. Rabbit anti-E1A and mouse anti-MYC are homemade antibodies. Mouse monoclonal anti-P21 was from Pharmingem™ (SX118).

### Xenograft

IMR90 and PC3 cells expressing FOXK1, FOXK2 or FOXK1 mutants were transduced with different combinations of oncogenes by retroviral transduction to evaluate their oncogenic transformation ability as previously established^52^. Transformed cells were trypsinized, counted and then resuspended in culture media supplemented with an equivalent volume of Matrigel^®^ (Corning™, 356237). About 2 × 10^6^ cells were subcutaneously injected (0.1 ml) using a 21-gauge needle in the right and left flank of each 6-week aged athymic nude mice (JAX002019, Jackson Laboratory^®^). Tumor size was followed at several points post injection by measuring the length and width of the tumor using a caliper. Tumor volume was calculated with the following formula (4/3*(3.14159)*(Length/2)*(Width/2)^2). All xenograft experiments were performed on both male and female individuals except for PC3 prostate cancer cells which were performed on male mice only. Tumor penetrance was calculated as a percentage of tumors observed compared to the total number of engraftments. Tumor presence was defined as tumor size of at least 0.1 cm^3^. Tumor latency was defined as mean time until tumor reached 0.1 cm^3^. All animal studies were approved by the Animal Care Committee of the research center of the Maisonneuve-Rosemont Hospital and in agreement with the guidelines of the Canadian Council on Animal Care.

### Retroviral transduction

Retroviruses were produced in Phoenix-Ampho. Cells were plated in 15 cm tissue culture dishes, the next day cells were transfected at 70-80% confluence. For one dish, 30 µg of plasmid, 10 µg of pCMV-VSVG and 10 µg HELPER were mixed with 143 µl of 1 mg/ml PEI (Sigma, 408727), incubated for 45 min, and then added to the cells. The cell media was changed 16 hours post-transfection, and retrovirus containing supernatants were collected at 48, 72 and 96 hours post-transfection. The viral supernatant was filtered through 0.45 µm filter and added to the target cells along with 8 µg/ml polybrene (Sigma, H9268). Following one to three infections, 16 hours each, the cells were selected with 2 µg/ml puromycin (Bioshop, PUR333) for 48 hours.

### Mass Spectrometry

Immuno-purified FOXK1 protein was subjected to SDS-PAGE and protein bands were stained with Coomassie brilliant blue. Following gel-extraction, reduction of samples was performed by adding 5 mM DTT in 50 mM ammonium bicarbonate. Alkylation was performed with 50 mM chloroacetamide and 50 mM ammonium bicarbonate. Trypsin digestion was performed for 8 hours at 37°C. Samples were loaded and separated on a homemade reversed-phase column (150 µm i.d. x 150 mm) with a 106-min gradient from 0–40% acetonitrile (0.2% FA) and a 600 nl/min flow rate on an Easy nLC-1,000 (Thermo Fisher Scientific) connected to an LTQ-Orbitrap Fusion (Thermo Fisher Scientific). Each full MS spectrum acquired with a 70,000 resolution was followed by 10 MS/MS spectra, where the 10 most abundant multiply charged ions were selected for MS/MS sequencing. Tandem MS experiments were performed using high-energy C-trap dissociation (HCD) and electron transfer dissociation (ETD) acquired in the Orbitrap. Peaks were identified using a Peaks 7.0 (Bioinformatics Solution Inc.) and peptide sequences were blasted against the human Uniprot database (74,530 sequences). Tolerance was set at 10 ppm for precursor and 0.01 Da for fragment ions during data processing and with carbamidomethylation (C), oxidation (M), deamidation (NQ), and Hex-N-acylation (ST) as variable modifications.

### Plasmid and siRNA transfections

For protein expression, HEK293T and HeLa cells were transfected with the mammalian expressing vectors using PEI. Three days post-transfection, cells were harvested for immunoblotting or immunoprecipitation. For RNAi-mediated protein depletion, IMR90 or U2OS cells were transfected twice with siRNA at 24h interval. Transfections were done with 150 pmol of siRNA in complete DMEM medium for 8-10 hours using Lipofectamine™ RNAi max (ThermoFisher Scientific, 13778150). Media was changed following transfection incubation. A pool of non-target siRNAs were used as a control. FOXK1 siRNAs (SASI_Hs01_00149056: GAUUGUAUGAUUCUGGGAA) and (SASI_Hs01_00149058: CUCUCUUUGAACCGUUACU) were obtained from Sigma-Aldrich.

### Native immunoprecipitation

Cells were lysed for 30 minutes on ice in EB300 buffer (50 mM Tris HCl pH 7.5, 300 mM NaCl, 5 mM EDTA, 0.5 % Triton, 1 mM DTT, 1 mM PMSF (Bioshop, PMS123100), 10 µM PUGNAc, 10 mM β-Glycerophosphate (Bioshop, GYP001), 1 mM Na_3_VO_4_ (Sigma, S6508), 50 mM NaF (Sigma, S7920) and 1x anti-protease cocktail inhibitors (Sigma, P8340). Lysates were centrifuged for 20 minutes at 14,000 rpm at 4 °C to pellet insoluble material and chromatin. Supernatants were adjusted to a final concentration of 150 mM NaCl and then incubated overnight with rotation at 4°C with either anti-Flag beads (Sigma, A2220) or with 1-5 μg of the appropriate antibody and then with Protein G Sepharose beads (Sigma, P3296). The following day, beads were washed with EB150 buffer (50 mM Tris pH 7.5, 150 mM NaCl, 5 mM EDTA, 0.5% Triton, 1 mM DTT, 1 mM PMSF, 2 µM PUGNAc, 10 mM β-Glycerophosphate, 1 mM Na_3_VO_4_, 50 mM NaF,1X anti-protease). For anti-Flag immunoprecipitation, bound proteins were eluted three times, 2 hours each, with 200 µg/ml of Flag peptide. The eluted material was supplemented with 2X Laemmli buffer and used for immunoblotting. For immunoprecipitation of endogenous proteins, protein G Sepharose beads were directly resuspended in 2X Laemmli buffer and subjected to immunoblotting.

### Immunoprecipitation under denaturing conditions

Cells were harvested in PBS and the pellet lysed in 25 mM Tris-HCl pH 7.3 containing 1% SDS. Following heating at 95 °C for 10 min, the cell extract is diluted ten times in 50 mM Tris-HCl pH7.5, 150 mM NaCl, 5 mM EDTA, 1% Triton X-100, 1X anti-protease, 1 mM PMSF, 1 mM DTT and 2 µM PUGNAc. The sample was centrifuged at 14,000 rpm for 20 min at 4°C and incubated with anti-FOXK1 antibody or anti-Flag beads overnight. Following pulldown with protein G-agarose beads or anti-Flag beads, FOXK1 is eluted with 2X Laemmli buffer or 200 µg/ml of Flag peptide diluted in 50 mM Tris-HCl pH7.5, 50 mM NaCl, 5 mM EDTA, 0.1% NP40, 1X anti-protease, 1 mM PMSF, 1 mM DTT and subsequently used for immunoblotting.

### Immunofluorescence

Cells were fixed in PBS containing 3 % PFA for 20 min. For antigen retrieval, the samples were incubated in sodium citrate Buffer (10 mM sodium citrate, 0.05% Tween 20, pH 6.0) and heated for 30 s in the microwave. The cells were then washed three times and permeabilized by incubation for 30 min in PBS containing 0.5% Triton X-100. Non-specific sites were blocked for 1 hour using PBS containing 0.1% Triton X-100 and 10% NBS. The coverslips were then incubated with primary antibodies for 3 hours at room temperature or overnight at 4 °C. After three washes of 15 min each, cells were incubated with secondary anti-mouse Alexa Fluor^®^ 594 (1/1,000) or anti-mouse Alexa Fluor^®^ 488 (1/1,000) and anti-rabbit Alexa fluor^®^ 488 (1/1,000) or anti-rabbit Alexa Fluor Alexa Fluor^®^ 488 594 (1/1,000) antibodies for 1 hour. Nuclei were stained with 4′, 6-diamidino-2-phenylindole (DAPI) during incubation with secondary antibodies. The images were acquired using DeltaVision Elite system (GE Healthcare) with z-stacking. Gamma, brightness, and contrast were adjusted on displayed images using the CellSens software. The collected EPI-fluorescence images were processed using ImageJ^53^ and Fiji^54^.

### Recombinant protein purification

BL21 CodonPlus-RIL bacteria were obtained from Agilent (230240) and were transformed with plasmids to produce GST-or His-tagged recombinant proteins. Cells were grown at 37°C and then treated with 400 µM Isopropyl β-d-1-thiogalactopyranoside (IPTG, Biobasic, IB0168) to induce protein expression. Cells were lysed on ice in 50 mM Tris-HCl pH 7.5, 100 mM NaCl, 1 mM EDTA, 1 % NP40, 1 mM PMSF, 0.5 mM DTT, and 1x anti-protease and sonicated. Cell lysates were incubated with GSH beads (Sigma, G4510) at 4°C for 5 hours. The beads were subsequently washed 6 times and an aliquot was used to assess the protein purification quality by SDS-PAGE. For His-OGT purification, bacteria were lysed in 50 mM Tris-HCl pH 8.0, 500 mM NaCl, 3 mM β-mercaptoethanol, 1 mM PMSF and 1× anti-protease. Following sonication, the bacteria lysates were incubated with Ni-NTA Agarose resin (Invitrogen^®^, R901-15) overnight at 4 °C. The resin was subsequently washed 5 times with 20 volumes of 50 mM Tris-HCl pH 8.0, 500 mM NaCl, 3 mM β-mercaptoethanol, 1 mM PSMF, 1× anti-protease, 20 mM imidazole. Proteins were eluted 3 times with 50 mM Tris-HCl pH 8.0, 500 mM NaCl and 200 mM imidazole, 3 mM β-mercaptoethanol and 1 mM EDTA. Arginine (200 µg/ml) was added to the elution buffer to prevent OGT precipitation. Protein eluates was then used for Flag affinity purification following the same procedure as for immunoprecipitation.

### GST-pulldowns

Recombinant GST-FOXK1 or its corresponding GST-FOXK1 protein fragments were kept immobilized on glutathione agarose beads. About 3 to 5 μg of bound proteins were incubated with the same quantity of His-OGT for 6 hours at 4 °C in GST pull down buffer containing 50 mM Tris-HCl, pH 7.5; 50 mM NaCl; 0.02% Tween 20; 500 μM DTT and 1 mM PMSF. The beads were washed 5 times with the same buffer, and FOXK1 bound proteins were eluted in 2X Laemmli buffer and subjected to Coomassie blue staining or immunoblotting.

### Immunodepletion of FOXK1 and Wheat Germ Agglutinin (WGA) purification

IMR90 cells were harvested in PBS and the cell pellet lysed in 25 mM Tris-HCl pH 7.3 containing 1% SDS. Following heating at 95 °C for 10 min, the cell extract is diluted in 50 mM Tris-HCl pH7.5, 150 mM NaCl, 5 mM EDTA, 1% Triton X-100, 1X anti-protease, 1 mM PMSF, 1 mM DTT and 2 µM PUGNAc. The sample was centrifuged at 14,000 rpm for 5 min at 4°C and incubated with anti-FOXK1 antibody overnight. Following pulldown with protein G-agarose beads, FOXK1 is eluted with 1 % SDS and the resulting material is diluted in 50 mM Tris-HCl pH7.5, 150 mM NaCl, 5 mM EDTA, 1% Triton X-100, 1X anti-protease, 1 mM PMSF, 1 mM DTT and 2 µM PUGNAc, before loading on WGA lectin resin (Vector Laboratories, #AL-1023). Following 6 hours incubation at 4 °C, several washes with the same buffer, FOXK1 is eluted with 2X Laemmli buffer. All fractions including inputs, washes and elutions were subjected to immunoblotting.

### OGT activity assay

In vitro O-GlcNAcylation reaction was conducted in 50 mM Tris-HCl, pH 7.5, containing 5 mM MgCl_2_, 1 mM of UDP-GlcNAc (Sigma, A8625) and 3 or 5 µg of purified His-OGT and mixed with either 3 or 5 µg of GST-FOXK1, GST-FOXK1 fragments or GST-FOXK2 bounds to beads. Purified GST was used as a control. The enzymatic assay was performed for the indicated times at 37°C and the reaction was stopped with 2X Laemmli buffer. Protein O-GlcNAcylation level was detected by immunoblotting.

### Immunoblotting

Cells were washed with PBS and lysed in 25 mM Tris-HCl pH 7.3 containing 2% SDS. Whole cell lysates were heated at 95°C for 5 min and sonicated. Protein quantification was done by bicinchoninic acid (BCA, Pierce™, 23222) assay and samples were diluted in Laemmli buffer. Proteins were resolved on 8 %, 10 % or 15 % Bis-Tris acrylamide gels and transferred to PVDF membrane, blocked for 1 hour in PBS containing 5 % non-fat milk, 0.1 % Tween-20, 5 mM sodium azide and 250 µg/ml Kanamycin (PBS-MT). Membranes were incubated 3 hours at room temperature or overnight at 4 °C with relevant primary antibodies (diluted in PBS-T containing 1% BSA, 5 mM sodium Azide and 250 µg/ml Kanamycin), subsequently washed 3 times in PBS-T and incubated for 1 hour with HRP-labeled secondary antibodies (diluted 1/1,000 in PBS-T containing 1% BSA and 250 µg/ml Kanamycin). Membranes were then washed three times in PBS-T. The band signals were acquired using an Azure C600 camera (Azure biosystem) and processed with Adobe Photoshop.

### qRT-PCR

Total RNA extracts were prepared using TRIzol™ (Invitrogen, 155960189) according to the manufacturer’s protocol. Total RNA (2 μg) was reverse transcribed in a final volume of 10 μL using SuperScript™ III Reverse Transcriptase kit (ThermoFisher Scientific^®^,) with oligo-p(dt)15 (Roche^®^, 10814270001). The cDNAs were analyzed by real time PCR using SYBR Green (Bimake, 21203) DNA quantification kit. The Applied Biosystems 7500 Real-Time PCR System (ThermoFisher Scientific) was used to detect the amplification levels and was programmed with an initial step of 3 min at 95°C, followed by 40 cycles of: 5 s at 95°C and 30 s at 60°C. All reactions were run in triplicate and the average values of Cts were used for quantification. The relative quantification of target genes was determined using the ΔΔCT method. The following primers were used: E2F1-F: AGACCCTGCAGAGCAGATGGTTAT, E2F1-R: TCGATCGGGCCTTGTTTGCTCTTA, Cyclin A-F: GCTGGAGCTGCCTTTCATTTAGCA, Cyclin A-R: TTGACTGTTGTGCATGCTGTGGTG, p21-F: CTGTCACTGTCTTGTACCCTTG, p21-R: CTTCCAGGACTGCAGGCTTCCTG, CDC6-F: GGAAGCCTTTACCTTTCTGGTG, CDC6-F: CAGCTGGCCTGGATACCTCTTC, MCM3-F: TGGGGATTCATACGACCCCT, MCM3-R: GAACACATCCAAGAGGGCCA, Actin-F: GAGCACAGAGCCTCGCCTTTG, Actin-R: CGAAGCCGGCCTTGCACATGC.

### ChIP-seq

Culture dishes containing 60 million cells were used per condition. Cells were fixed in culture media for 10 minutes in 1% formaldehyde (F1635, Sigma) at room temperature. Cells were quenched for 5 minutes with 125 mM L-glycine in ice cold PBS and quickly washed with ice cold PBS. Cells were lysed in 0.25% Triton X-100, 10 mM Tris pH8, 10 mM EDTA and 0.5 mM EGTA with anti-protease for 5 min on ice. Cells were then resuspended in 200 mM NaCl, 10 mM Tris pH8, 1 mM EDTA, 0.5 mM EGTA with anti-protease and incubated for 30 min on ice. Cells were split in 3 tubes and sonicated on ice in 10 mM Tris pH 8 containing 0.5% SDS, 0.5% Triton X-100, 0.05% sodium deoxycholate, 140 mM NaCl, 1 mM EDTA, 1 mM EGTA and anti-protease to yield mean fragment size of 500 bp. The chromatin suspension was centrifuged at 14,000 rpm for 15 minutes at 4°C and the supernatant incubated overnight with pre-coupled Dynabeads^®^ (G + A, ThermoFisher Scientific, 10002D) with anti-Flag M2 antibody (F3165, Sigma) or an antibody targeting a protein of interest. About 50 μL of chromatin was kept as an input. The beads were then washed successively at room temperature in low salt buffer (1 % Triton X-100, 0.1% SDS, 150 mM NaCl, 20 mM Tris pH 8, 2 mM EDTA), high salt buffer (1 % Triton X-100, 0.1% SDS, 500 mM NaCl, 20 mM Tris pH 8, 2 mM EDTA), LiCl buffer (1 % NP40, 250 mM LiCl, 10 mM Tris pH 8, 1 mM EDTA) and TEN buffer (50 mM NaCl, 10 mM Tris pH 8, 1 mM EDTA). Beads and inputs were then decrosslinked overnight at 65°C in 1% SDS, 50 mM Tris pH 8, 10 mM EDTA. Samples were treated with RNAse (100 μg/ml) at room temperature for 15 min and then with proteinase K (825 μg/ml, NEB™, P8107S) for 1 hour at 65°C. Samples were purified using a DNA purification micro column (QIAquick™ PCR purification kit, QIAgene^®^, 28106). The inputs were first treated with phenol chloroform and DNA was precipitated by adding sodium acetate. Once resuspended, the input DNA was also purified using the DNA purification micro column. At least six independent ChIPs were pooled before library preparation. The preparation of next generation sequencing libraries was done using the KAPA HyperPrep ChIP Library kit (Roche Sequencing solutions) at the molecular biology and functional genomics platform of the Institut de Recherche Clinique de Montréal (IRCM). The ChIP libraries were sequenced on an Illumina Novaseq 6000 sequencer with a sequencing depth of 50 million reads minimum per condition (service provided by Genome Quebec).

### Cut&Run

Cut&Run assays were performed in accordance with the manufacturer’s recommendations, with specific modifications, using the CUTANA™ ChIC/CUT&RUN Kit (Epicypher^®^, 14-1048). IMR90 cells were washed once with PBS and scraped. Cellular nuclei were extracted on ice using 20 mM HEPES (pH 7.9) containing 10 mM KCl, 0.1% Triton X-100, 20% glycerol, and 1 mM MnCl_2_, supplemented with 0.5 mM spermidine and 1X complete™ Mini EDTA-free Protease Inhibitor Cocktail (Roche, 11836170001). The nuclei were then further processed with a loose-fit Dounce homogenizer. A total of 500,000 nuclei were bound to concanavalin A magnetic beads (ConA) for 10 minutes at room temperature. The beads-nuclei mixtures were subsequently resuspended in 140 µl of antibody buffer, consisting of 20 mM HEPES (pH 7.9), 150 mM NaCl, 0.01% digitonin, 1 mM EDTA, and 1 X anti-protease, 0.5 mM spermidine, and 10 µM PUGNAc. Antibody incubation was carried out at 4°C for 2 hours. The ConA beads were then resuspended in 100 µl of 20 mM HEPES pH 7.9, 150 mM NaCl, 0.01% digitonin, 1X anti-protease, 0.5 mM spermidine, and 10 µM PUGNAc and treated with MNase at 4°C for 2 hours. The reaction was terminated by adding 66 µl of stop buffer (340 mM NaCl, 20 mM EDTA, 4 mM EGTA, 50 µg/ml RNase A, 50 µg/ml glycogen and spike-in *E. coli* DNA). Library preparation was performed at the Molecular Biology and Functional Genomics Core Facility of IRCM. We used 1 µg of rabbit polyclonal anti-BAP1 (Cell Signaling, D1W9B). For H2AK119ub Cut&Run, MNase digestion was carried out for 30 minutes and 1 µg of rabbit anti-H2AK119ub was used (Cell Signaling, D27C4). For H3K4me1 Cut&Run, we used 1 µg of rabbit anti-H3K4me1 (Abcam, 8895). The DNA was subsequently purified and the libraries were prepared and sequenced on an Illumina NovaSeq 6000 instrument. The sequencing depth was 10 million reads par condition except H2AK119ub, 50 million reads were acquired.

### ATAC-seq

We counted cells and extracted 50,000 nuclei per condition by incubating cells 30 min at 4°C in hypotonic cell lysis buffer containing sodium citrate tribasic dihydrate (0.1% (wt/vol) and 0.1% (vol/vol) Triton X-100. Nuclei were then resuspended in normal cell lysis buffer (10 mM Tris-HCl, pH 7.4, 10 mM NaCl, 3 mM MgCl_2_ and 0.1% (vol/vol) IGEPAL CA-630) for 30 min at 4°C. Transposition was performed directly on nuclei following manufacturer recommendations (Tn5 Illumina) at the molecular biology and functional genomics platform of the Institut de Recherche Clinique de Montréal (IRCM). DNA was then purified and enriched by PCR, and the library was recovered with GeneRead Purification columns (QIAgen^®^) and sequenced on an Illumina NovaSeq 6000 instrument.

### RNA-seq

A biological triplicate was harvested 5 days following IMR90 retroviral infection in TRIzol reagent. Total RNAs were extracted by phenol/chloroform treatment followed by additional purification on column (RNeasy^®^ Mini Kit, QIAgene, 74204) following manufacturer protocol. The libraries were prepared at the molecular biology and functional genomic platform of the IRCM using ribosomal RNA depletion (KAPA RNA HyperPrep kit) with a sequencing depth of minimum 50 million reads on an Illumina NovaSeq 6000 instrument.

### Quantification and Statistical Analysis

For genome occupancy studies, we mapped ChIP–seq, Cut&Run, and ATAC–seq reads on the human genome assembly GRCh38 by using bowtie2 v2.3.1 with the following settings: -p –fr --no- mixed --no-unal -x -1 -2 -S^55^. Optical and PCR duplicates reads were removed using picard v2.9.0 (https://broadinstitute.github.io/picard/). We processed the mapped sequence reads with MACS2^56^ version 2.1.1 using the parameters -t -c -n --outdir -f BAMPE -p 1e-7 -g --call-summits 0.00001. Peaks annotation and motif analysis was performed with HOMER^57^ using default setting. For motif analysis, we use –len parameter with length of 6, 8, 10, 12, 14, 16. We used deeptools^58^ to generate heatmap and plot profile of ChIP-seq and ATAC-seq and Cut&Run. Highly correlated replicates samples from Cut&Run experiments were merged for further visualization.

For RNA-seq experiments the quality of the raw reads was assessed with FASTQC v0.11.8 (https://www.bioinformatics.babraham.ac.uk/projects/fastqc/). After examining the quality of the raw reads, no trimming was deemed necessary. The human samples were spiked-in with drosophila S2 cells. A hybrid reference genome and annotation concatenating both species was used for the alignment. The GRCh38 (release 102) reference genome and BDGP6.32 (release 107) reference genome were used and downloaded from Ensembl^59^. The reads were aligned to the hybrid reference genome with STAR v2.7.6a^60^. The raw counts were calculated with FeatureCounts v1.6.0^61^ based on the hybrid reference genome. Differential expression was performed using DESeq2 v1.30.0 R package^62^. Differentially expressed genes (DEGs) heatmap was drawn based on z-score of normalized count. The ontology analysis was performed on the significant DEGs using Enrichr^63–65^. Odds ratio takes into account the number of overlapping genes with the annotated set (a), the size of the annotation set (b), the total number of genes in the input (c) and the total number of genes in the human genome (d). The computation is as follows: oddsRatio = (1* a * d) / Math.max(1 * b * c, 1).

The GSEA analysis was performed on the hallmark gene sets collection with all normalized counts from DESeq2 using gseapy v1.0.0 python package^66^. The MAplot shows the distribution of the differences of mean expression given by DESeq2 between 2 samples. Figures were generated using R language (https://www.R-project.org/) and python language (http://www.python.org).

### Gene expression analysis in cancer

Gene expression profiles in human cancerous and normal tissues (TCGA, TARGET and GTEx datasets) were obtained from UCSC Xena (https://xenabrowser.net/datapages/)^67^. Data from cell lines were removed from subsequent analysis. In each tissue, Pearson correlation between FOXK1 and FOXK2 expression and its statistical significance were calculated in cor.test function in R (v4.0.5) (https://www.R-project.org/). For read counts of FOXK1 and FOXK2, we retrieve TCGA cancer samples transcript counts using TCGAbiolinks^68^ R package. Normalized reads counts were then sorted between the top 10% and bottom 10% expression of FOXK1 and FOXK2. Read counts were transform to z-score for visualization.

## Acknowledgments

We thank the Dre Virginie Calderon and Gaspard Reulet from the IRCM genomics and bioinformatics for their helps with bioinformatics analysis. We thank Dr David Vocaldo for generously providing OGT and OGA inhibitors. We thank, Jihane Khalifa, Chloé Ducquette, Bintou Haidara, Maxime Uriarte, Salima Daou, Malik Affar and Diana Adjaoud for technical assistance. We also thank the technical assistants of the animal care, bio-imaging and flow cytometry facilities of CR-HMR. This work was supported by grants from the Canadian Institutes of Health Research (CIHR) (MOP159539) to E.B.A. Molecular graphics and analyses performed with UCSF ChimeraX, developed by the Resource for Biocomputing, Visualization, and Informatics at the University of California, San Francisco, with support from National Institutes of Health R01-GM129325 and the Office of Cyber Infrastructure and Computational Biology, National Institute of Allergy and Infectious Diseases. The author generated some code for data visualization with GPT-4, OpenAI’s large-scale language-generation mode (https://chat.openai.com/chat). Upon generating code, the author reviewed, edited, and revised the code to their own liking and takes ultimate responsibility for the content of this publication.

## Author Contribution

LM, OA, EBA contributed study design. MGL, KB, BE, ME, MP, AB, SM, AS, AB, LH, YT, FAM contributed to experiments, data Collection, and data Analysis. LM, OA, EBA, wrote the original manuscript Writing. AB, EB, PT, JD contributed additional help with Design and Data Analysis. All authors contributed to critical revisions, final editing and corrections of the manuscript.

## Extended Data legends

**Extended Data Figure 1:**

**FOXK1 overexpression in cancer is associated with a poor prognosis**

**a**-**b**) Comparison of FOXK1 and FOXK2 expression between cancer and normal tissues. Cancer data were retrieved from TCGA TARGET GTEx dataset. P-value is calculated by Wilcoxson test. **c**) Comparison of co-expression of FOXK1, FOXK2 and other FOX genes between cancer and normal tissues. FOX genes were sorted by Pearson correlation with FOXK1 and FOKX2. **d**) Kaplan-Meier survival curve of patients from the TCGA database presenting high or low mRNA levels of FOXK1 or FOXK2 in cancer tissues. **e**) Proliferation of U2OS and HCT116 cells stably expressing FOXK1, FOXK2 or empty vector was analyzed by colony forming assay (CFA). Crystal violet signal for each condition was quantified using ImageJ and plotted in the right. Results from one representative experiment are shown.

**Extended Data Figure 2:**

**FOXK1 and FOXK2 regulate overlapping and specific gene expression programs**

**a**) Transcript counts of FOXK1 and FOXK2 from our RNA-seq experiment in IMR90 cells expressing either empty vector, FOXK1 or FOXK2. Mean transcript counts for each condition are represented in the adjacent table. **b**) Transcript counts from TCGA cancer samples were retrieved and categorized into two groups: the top 10% with the highest expression (mean FOXK1 = 3193, FOXK2 = 3115) and the top 10% with the lowest expression (mean FOXK1 = 398, FOXK2 = 757) of FOXK1 and FOXK2 transcripts. Counts for FOXK1 and FOXK2 in both group were plotted as boxplots. **c**) Heat map representing the transcript count (z-score) of genes differentially expressed between FOXK1 and FOXK2. GO analysis (MSigDB hallmark) was performed for each gene cluster. **d**) MA Plot representing the mean expression against the log fold change of genes when comparing IMR90 cells overexpressing FOXK2 with cells expressing the empty vector. **e**) GO analysis performed on genes differentially regulated between FOXK2 and control conditions. **f**) GSEA performed on genes deregulated (log fold change greater than 0.6) between conditions of FOXK1 overexpression and empty vector or FOXK2 overexpression and empty vector. Enrichment of genes associated to autophagy and DNA replication are represented.

**Extended Data Figure 3:**

**FOXK1 genome occupancy of cell cycle genes**

**a**) Correlation plot between endogenous and exogenous (Flag-tagged) ChIP-seq peaks of FOXK1 and FOXK2 in IMR90 cells. **b**) GO analysis of common promoters targeted by endogenous or exogenous FOXK1 in IMR90, U2OS and K562 as determined by ChIP-seq. **c**) GO analysis of genes up-regulated by FOXK1 in IMR90 found in the cell cycle pathway (KEGG) containing a ChIP-seq signal of FOXK1 in their promoters (red star).

**Extended Data Figure 4:**

**FOXK1 O-GlcNAcylation during cell cycle progression and cell differentiation**

**a**) Immunoprecipitation of endogenous FOXK1 and analysis of its O-GlcNAc levels in murine 3T3L1 cells treated with the OGA inhibitor, PUGNAc, or OGT inhibitor (OGTi). **b**) FOXK1 cellular localization following treatment with OGA inhibitor, OSMI-4, or with OGT inhibitor, PUGNAc, was analyzed by immunofluorescence in IMR90 cells. The non-relevant USP10 protein serves as a control for the cytoplasmic compartment. Representative of three independent experiments. **c**) U2OS cells were deprived of serum for 24h to synchronize cells in G1 phase. Cells were then stimulated with the addition of serum and FOXK1 was immunoprecipitated at different times to analyze its O-GlcNAcylation levels. CDC6 was used as a control of cell synchronization. **d**) Human lung fibroblasts (HLF) were synchronized by contact inhibition for several days to induce G0 entry. Cell cycle block release was performed by trypsinization and plating at low density in fresh medium. FOXK1 was immunoprecipitated at different times to analyze its O-GlcNAcylation. CDC6 was used as a control of synchronization. **e**) Pre-adipocytes 3T3-L1 were differentiated into adipocytes and FOXK1 was immunoprecipitated to analyze its O-GlcNAcylation levels upon differentiation. Fabp4 and Perilipin are markers of adipocytes.

**Extended Data Figure 5:**

**Mapping of FOXK1 region and sites targeted by O-GlcNAcylation**

**A**) Recombinant GST-FOXK1 fragments pulldown with OGT to determine its interaction motif with FOXK1. Representative of two experiments. **B**) Mass spectra of FOXK1 residues targeted by O-GlcNAcylation.

**Extended Data Figure 6:**

**Characterization of FOXK1 O-GlcNAcylation sites and impact of O-GlcNAcylation on protein stability and localization.**

**a**) PC3 **b**) U2OS and **c**) K562 cells stably expressing Flag tagged version of FOXK1, FOXK1^7A^ or FOXK1^11A^ were harvested for Flag immunoprecipitation and O-GlcNAcylation detection. The star (*) represent non-specific bands in O-GlcNAc signal from U2OS cells. **d**) Transient transfection in HeLa cells of individual O-GlcNAc-modified residues mutation in FOXK1 (Flag tagged) to assess their O-GlcNAcylation levels after immunoprecipitation. The O-GlcNAc levels of FOXK1 mutants was quantified using ImageJ and plotted (right histogram) (n=3). **e**) Transient transfection in HeLa cells of combined O-GlcNAc-modified residues mutation in FOXK1 to assess O-GlcNAcylation levels following immunoprecipitation. The O-GlcNAc levels of FOXK1 mutants was quantified using ImageJ and plotted (right histogram) (n=4). **f**) U2OS cells overexpressing FOXK1, FOXK1^7A^ or FOXK1^11A^ were treated with 20 μg/ml cycloheximide or 20 μM MG132 and harvested for protein levels assessment by immunoblotting. CDC6 was used as a control for treatment efficacy. Flag-FOXK1 signal was quantified and normalized to tubulin signal. Representative of three independent experiments. **G**) Sub-cellular localization of exogenous FOXK1, FOXK1^7A^, FOXK1^11A^ and FOXK2 in U2OS cells. Detection of HSP90⍺/β was used as a control of the cytoplasm compartment. Representative of three independent experiments.

**Extended Data Figure 7:**

**Effect of FOXK1 O-GlcNAcylation on tumor formation and progression**

**a**) Western blot of IMR90 expressing empty vector, FOXK1, FOXK1^7A^ or FOXK1^11A^ in combination with E1A and RAS^G12V^. The full combination containing HDM2 + E1A + RAS^G12V^ is also shown. **b**) Left: IMR90 tumors expressing HDM2 + RAS^G12V^ in combination with FOXK1 or FOXK1^11A^ at the time of harvest. Right: Graph representing final volume of tumors expressing FOXK1 or FOXK1^11A^ in combination with HDM2 + RAS^G12V^, and in comparison with tumors expressing HDM2 + E1A + RAS^G12V^. **c**) Left: IMR90 tumors expressing FOXK1, FOXK1^7A^ or FOXK1^11A^ with E1A (minimal combination) at the time of harvest. Right: Graph of final size of tumors expressing FOXK1, FOXK1^7A^ or FOXK1^11A^ in combination with E1A, and in comparison with tumors expressing HDM2 + E1A + RAS^G12V^. **d**) U2OS cells stably expressing empty vector, siRNA-resistant FOXK1 cDNA (FOXK1 or FOXK1^7A^) were transfected with siRNA non-target (NT) or siRNA targeting endogenous FOXK1. Left: western-blotting depicting FOXK1 or FOXK1^7A^ expression and the efficiency of endogenous FOXK1 depletion by siRNA. Middle: Cells were plated to perform colony forming ability (CFA). Right: Violet crystal was extracted from cells and intensity was quantified by absorbance (technical triplicates). **e**) mRNA expression of Cyclin A in U2OS expressing FOXK1 or FOXK1^7A^ . Biological replicates. **f**) PC3 stably expressing empty vector, FOXK1, FOXK1^7A^ or FOXK1^11A^ were plated at low density for several days. Cells were stained with crystal violet. **g**) PC3 expressing empty vector, FOXK1, FOXK1^7A^ or FOXK1^11A^ were engrafted subcutaneously in the flanks of nude mice. Mice were sacrificed once the tumors reached the limit point. **h**) Images of tumors at the time of harvest. **i**) E2F1 protein levels were analyzed by western blotting on cell extracts of PC3 expressing empty vector, FOXK1, FOXK1^7A^ or FOXK1^11A^. Data are represented as mean ± SEM. One-way ANOVA with Dunnett’s multiple comparisons was used (**b**, **c**, **d**, **g**). Statistical t-test (**e**) \**P* < 0.05, \*\**P* < 0.01, \*\*\*\**P* < 0.0001.

**Extended Data Figure 8:**

**Effect of FOXK1 O-GlcNAcylation on genomic FOXK1 and BAP1 occupancy**

**a**) Correlation plot between exogenous ChIP-seq Flag signal for FOXK1, FOXK1^7A^ and FOXK1^11A^ in IMR90 cells. **b**) Chromatin occupancy of exogenous Flag tagged FOXK1, FOXK1^7A^ and FOXK1^11A^ in IMR90 cells on distal regions. Distal regions, corresponding to regions containing FOXK1 binding 1kb away upstream and downstream from TSS. **c**) Co-localization between FOXK1 and FOXK1^7A^ ChIP-seq with opened chromatin regions from ATAC-seq experiments performed in U2OS cells overexpressing siRNA resistant cDNA of FOXK1 or FOXK1^7A^. U2OS cells were treated with siRNA targeting FOXK1 for 72h before performing ATAC-seq experiment. ChIP-seq and ATAC-seq signals are centered on regions containing FOXK1 peaks. **d**) Occupancy of FOXK1, FOXK1^7A^ and FOXK1^11A^ assessed by ChIP-seq of 3-Flag tagged proteins in IMR90 on promoters and surrounding distal regions of genes whose expression is associated with FOXK1 overexpression.

**Extended Data Figure 9:**

**Effect of FOXK1 O-GlcNAcylation on epigenomic histone marks**

**a**) Analysis of BAP1 recruitment by Cut&Run in IMR90 cells expressing FOXK1, FOXK1^7A^ or FOXK1^11A^ on promoters of genes identified by RNA-seq as being differentially regulated by FOXK1 overexpression compared to FOXK2 or empty vector. Distal regions correspond to regions surrounding promoters at a distance greater than 1kb. These regions are enriched for H3K27Ac and BRD4 and were qualified as enhancers. **b**) Differential enrichment of H3K4me1 and H2AK119ub histone marks in IMR90 cells expressing FOXK1, FOXK1^7A^ or FOXK1^11A^ on the same promoters and distal regions.

## References

1 Matthews, H. K., Bertoli, C. & de Bruin, R. A. M. Cell cycle control in cancer. Nat Rev Mol Cell Biol 23, 74–88 (2022). 10.1038/s41580-021-00404-3

2 Kent, L. N. & Leone, G. The broken cycle: E2F dysfunction in cancer. Nat Rev Cancer 19, 326–338 (2019). 10.1038/s41568-019-0143-7

3 Sukonina, V. et al. FOXK1 and FOXK2 regulate aerobic glycolysis. Nature 566, 279–283 (2019). 10.1038/s41586-019-0900-5

4 He, L. et al. mTORC1 Promotes Metabolic Reprogramming by the Suppression of GSK3-Dependent Foxk1 Phosphorylation. Mol Cell 70, 949–960 e944 (2018). 10.1016/j.molcel.2018.04.024

5 Sakaguchi, M. et al. FoxK1 and FoxK2 in insulin regulation of cellular and mitochondrial metabolism. Nat Commun 10, 1582 (2019). 10.1038/s41467-019-09418-0

6 Bowman, C. J., Ayer, D. E. & Dynlacht, B. D. Foxk proteins repress the initiation of starvation-induced atrophy and autophagy programs. Nat Cell Biol 16, 1202–1214 (2014). 10.1038/ncb3062

7 Kang, Y., Zhang, K., Sun, L. & Zhang, Y. Regulation and roles of FOXK2 in cancer. Front Oncol 12, 967625 (2022). 10.3389/fonc.2022.967625

8 Shan, L. et al. FOXK2 Elicits Massive Transcription Repression and Suppresses the Hypoxic Response and Breast Cancer Carcinogenesis. Cancer Cell 30, 708–722 (2016). 10.1016/j.ccell.2016.09.010

9 Zhang, F. et al. FOXK2 suppresses the malignant phenotype and induces apoptosis through inhibition of EGFR in clear-cell renal cell carcinoma. Int J Cancer 142, 2543–2557 (2018). 10.1002/ijc.31278

10 Wang, B. et al. Forkhead box K2 inhibits the proliferation, migration, and invasion of human glioma cells and predicts a favorable prognosis. Onco Targets Ther 11, 1067–1075 (2018). 10.2147/OTT.S157126

11 Liu, X. et al. Downregulation of FOXK2 is associated with poor prognosis in patients with gastric cancer. Mol Med Rep 18, 4356–4364 (2018). 10.3892/mmr.2018.9466

12 Liu, Y. et al. FOXK2 transcription factor suppresses ERalpha-positive breast cancer cell growth through down-regulating the stability of ERalpha via mechanism involving BRCA1/BARD1. Sci Rep 5, 8796 (2015). 10.1038/srep08796

13 Chen, S. et al. Foxk2 inhibits non-small cell lung cancer epithelial-mesenchymal transition and proliferation through the repression of different key target genes. Oncol Rep 37, 2335–2347 (2017). 10.3892/or.2017.5461

14 Li, L., Gong, M., Zhao, Y., Zhao, X. & Li, Q. FOXK1 facilitates cell proliferation through regulating the expression of p21, and promotes metastasis in ovarian cancer. Oncotarget 8, 70441–70451 (2017). 10.18632/oncotarget.19713

15 Wu, Y. et al. Oncogene FOXK1 enhances invasion of colorectal carcinoma by inducing epithelial-mesenchymal transition. Oncotarget 7, 51150–51162 (2016). 10.18632/oncotarget.9457

16 Wu, M. et al. FOXK1 interaction with FHL2 promotes proliferation, invasion and metastasis in colorectal cancer. Oncogenesis 5, e271 (2016). 10.1038/oncsis.2016.68

17 Chen, D. et al. FOXK1 plays an oncogenic role in the development of esophageal cancer. Biochem Biophys Res Commun 494, 88–94 (2017). 10.1016/j.bbrc.2017.10.080

18 Zhang, H. et al. Coexpression of FOXK1 and vimentin promotes EMT, migration, and invasion in gastric cancer cells. J Mol Med (Berl) 97, 163–176 (2019). 10.1007/s00109-018-1720-z

19 Peng, Y. et al. Direct regulation of FOXK1 by C-jun promotes proliferation, invasion and metastasis in gastric cancer cells. Cell Death Dis 7, e2480 (2016). 10.1038/cddis.2016.225

20 Cao, H. et al. High FOXK1 expression correlates with poor outcomes in hepatocellular carcinoma and regulates stemness of hepatocellular carcinoma cells. Life Sci 228, 128–134 (2019). 10.1016/j.lfs.2019.04.068

21 Garry, D. J., Maeng, G. & Garry, M. G. Foxk1 regulates cancer progression. Ann Transl Med 8, 1041 (2020). 10.21037/atm-2020-94

22 Tsai, K. L. et al. Crystal structure of the human FOXK1a-DNA complex and its implications on the diverse binding specificity of winged helix/forkhead proteins. J Biol Chem 281, 17400–17409 (2006). 10.1074/jbc.M600478200

23 Durocher, D. et al. The molecular basis of FHA domain:phosphopeptide binding specificity and implications for phospho-dependent signaling mechanisms. Mol Cell 6, 1169–1182 (2000). 10.1016/s1097-2765(00)00114-3

24 Ferbeyre, G. et al. PML is induced by oncogenic ras and promotes premature senescence. Genes Dev 14, 2015–2027 (2000).

25 Pearson, M. et al. PML regulates p53 acetylation and premature senescence induced by oncogenic Ras. Nature 406, 207–210 (2000). 10.1038/35018127

26 Seger, Y. R. et al. Transformation of normal human cells in the absence of telomerase activation. Cancer Cell 2, 401–413 (2002). 10.1016/s1535-6108(02)00183-6

27 Mallette, F. A. & Richard, S. JMJD2A promotes cellular transformation by blocking cellular senescence through transcriptional repression of the tumor suppressor CHD5. Cell Rep 2, 1233–1243 (2012). 10.1016/j.celrep.2012.09.033

28 Serrano, M., Lin, A. W., McCurrach, M. E., Beach, D. & Lowe, S. W. Oncogenic ras provokes premature cell senescence associated with accumulation of p53 and p16INK4a. Cell 88, 593–602 (1997). 10.1016/s0092-8674(00)81902-9

29 Yu, H. et al. The ubiquitin carboxyl hydrolase BAP1 forms a ternary complex with YY1 and HCF-1 and is a critical regulator of gene expression. Mol Cell Biol 30, 5071–5085 (2010). 10.1128/MCB.00396-10

30 Machida, Y. J., Machida, Y., Vashisht, A. A., Wohlschlegel, J. A. & Dutta, A. The deubiquitinating enzyme BAP1 regulates cell growth via interaction with HCF-1. J Biol Chem 284, 34179–34188 (2009). 10.1074/jbc.M109.046755

31 Mashtalir, N. et al. Autodeubiquitination protects the tumor suppressor BAP1 from cytoplasmic sequestration mediated by the atypical ubiquitin ligase UBE2O. Mol Cell 54, 392–406 (2014). 10.1016/j.molcel.2014.03.002

32 Okino, Y., Machida, Y., Frankland-Searby, S. & Machida, Y. J. BRCA1-associated protein 1 (BAP1) deubiquitinase antagonizes the ubiquitin-mediated activation of FoxK2 target genes. J Biol Chem 290, 1580–1591 (2015). 10.1074/jbc.M114.609834

33 Daou, S. et al. Monoubiquitination of ASXLs controls the deubiquitinase activity of the tumor suppressor BAP1. Nat Commun 9, 4385 (2018). 10.1038/s41467-018-06854-2

34 Yang, X. & Qian, K. Protein O-GlcNAcylation: emerging mechanisms and functions. Nat Rev Mol Cell Biol 18, 452–465 (2017). 10.1038/nrm.2017.22

35 Saunders, H., Dias, W. B. & Slawson, C. Growing and dividing: how O-GlcNAcylation leads the way. J Biol Chem 299, 105330 (2023). 10.1016/j.jbc.2023.105330

36 Taparra, K. et al. O-GlcNAcylation is required for mutant KRAS-induced lung tumorigenesis. J Clin Invest 128, 4924–4937 (2018). 10.1172/JCI94844

37 Yi, W. et al. Phosphofructokinase 1 glycosylation regulates cell growth and metabolism. Science 337, 975–980 (2012). 10.1126/science.1222278

38 Lynch, T. P. et al. Critical role of O-Linked beta-N-acetylglucosamine transferase in prostate cancer invasion, angiogenesis, and metastasis. J Biol Chem 287, 11070–11081 (2012). 10.1074/jbc.M111.302547

39 Tasdemir, N. et al. BRD4 Connects Enhancer Remodeling to Senescence Immune Surveillance. Cancer Discov 6, 612–629 (2016). 10.1158/2159-8290.CD-16-0217

40 Wang, L. et al. Resetting the epigenetic balance of Polycomb and COMPASS function at enhancers for cancer therapy. Nat Med 24, 758–769 (2018). 10.1038/s41591-018-0034-6

41 Hu, D. et al. The MLL3/MLL4 branches of the COMPASS family function as major histone H3K4 monomethylases at enhancers. Mol Cell Biol 33, 4745–4754 (2013). 10.1128/MCB.01181-13

42 Hanahan, D. Hallmarks of Cancer: New Dimensions. Cancer Discov 12, 31–46 (2022). 10.1158/2159-8290.CD-21-1059

43 Slawson, C. & Hart, G. W. O-GlcNAc signalling: implications for cancer cell biology. Nat Rev Cancer 11, 678–684 (2011). 10.1038/nrc3114

44 Qin, J. et al. BAP1 promotes breast cancer cell proliferation and metastasis by deubiquitinating KLF5. Nat Commun 6, 8471 (2015). 10.1038/ncomms9471

45 Daou, S. et al. Crosstalk between O-GlcNAcylation and proteolytic cleavage regulates the host cell factor-1 maturation pathway. Proc Natl Acad Sci U S A 108, 2747–2752 (2011). 10.1073/pnas.1013822108

46 Narita, M. et al. A novel role for high-mobility group a proteins in cellular senescence and heterochromatin formation. Cell 126, 503–514 (2006). 10.1016/j.cell.2006.05.052

47 Jumper, J. et al. Highly accurate protein structure prediction with AlphaFold. Nature 596, 583–589 (2021). 10.1038/s41586-021-03819-2

48 Varadi, M. et al. AlphaFold Protein Structure Database: massively expanding the structural coverage of protein-sequence space with high-accuracy models. Nucleic Acids Res 50, D439–D444 (2022). 10.1093/nar/gkab1061

49 Pettersen, E. F. et al. UCSF ChimeraX: Structure visualization for researchers, educators, and developers. Protein Sci 30, 70–82 (2021). 10.1002/pro.3943

50 Mallette, F. A. et al. Transcriptome analysis and tumor suppressor requirements of STAT5-induced senescence. Ann N Y Acad Sci 1197, 142–151 (2010). 10.1111/j.1749-6632.2010.05192.x

51 Mallette, F. A., Gaumont-Leclerc, M. F. & Ferbeyre, G. The DNA damage signaling pathway is a critical mediator of oncogene-induced senescence. Genes Dev 21, 43–48 (2007). 10.1101/gad.1487307

52 Mason, D. X. et al. Defined genetic events associated with the spontaneous in vitro transformation of ElA/Ras-expressing human IMR90 fibroblasts. Carcinogenesis 27, 350–359 (2006). 10.1093/carcin/bgi264

53 Schneider, C. A., Rasband, W. S. & Eliceiri, K. W. NIH Image to ImageJ: 25 years of image analysis. Nat Methods 9, 671–675 (2012). 10.1038/nmeth.2089

54 Schindelin, J., et al. Fiji: an open-source platform for biological-image analysis. Nat Methods 9, 676-682 (2012). 10.1038/nmeth.2019

55 Langmead, B. & Salzberg, S. L. Fast gapped-read alignment with Bowtie 2. Nat Methods 9, 357–359 (2012). 10.1038/nmeth.1923

56 Zhang, Y. et al. Model-based analysis of ChIP-Seq (MACS). Genome Biol 9, R137 (2008). 10.1186/gb-2008-9-9-r137

57 Heinz, S. et al. Simple combinations of lineage-determining transcription factors prime cis-regulatory elements required for macrophage and B cell identities. Mol Cell 38, 576–589 (2010). 10.1016/j.molcel.2010.05.004

58 Ramirez, F. et al. deepTools2: a next generation web server for deep-sequencing data analysis. Nucleic Acids Res 44, W160–165 (2016). 10.1093/nar/gkw257

59 Martin, F. J. et al. Ensembl 2023. Nucleic Acids Res 51, D933–D941 (2023). 10.1093/nar/gkac958

60 Dobin, A. et al. STAR: ultrafast universal RNA-seq aligner. Bioinformatics 29, 15–21 (2013). 10.1093/bioinformatics/bts635

61 Liao, Y., Smyth, G. K. & Shi, W. featureCounts: an efficient general purpose program for assigning sequence reads to genomic features. Bioinformatics 30, 923–930 (2014). 10.1093/bioinformatics/btt656

62 Love, M. I., Huber, W. & Anders, S. Moderated estimation of fold change and dispersion for RNA-seq data with DESeq2. Genome Biol 15, 550 (2014). 10.1186/s13059-014-0550-8

63 Kuleshov, M. V. et al. Enrichr: a comprehensive gene set enrichment analysis web server 2016 update. Nucleic Acids Res 44, W90–97 (2016). 10.1093/nar/gkw377

64 Xie, Z. et al. Gene Set Knowledge Discovery with Enrichr. Curr Protoc 1, e90 (2021). 10.1002/cpz1.90

65 Chen, E. Y. et al. Enrichr: interactive and collaborative HTML5 gene list enrichment analysis tool. BMC Bioinformatics 14, 128 (2013). 10.1186/1471-2105-14-128

66 Fang, Z., Liu, X. & Peltz, G. GSEApy: a comprehensive package for performing gene set enrichment analysis in Python. Bioinformatics 39 (2023). 10.1093/bioinformatics/btac757

67 Goldman, M. J. et al. Visualizing and interpreting cancer genomics data via the Xena platform. Nat Biotechnol 38, 675–678 (2020). 10.1038/s41587-020-0546-8

68 Colaprico, A. et al. TCGAbiolinks: an R/Bioconductor package for integrative analysis of TCGA data. Nucleic Acids Res 44, e71 (2016). 10.1093/nar/gkv1507

