## Extended Figures for "O-GlcNAcylation of FOXK1 orchestrates the E2F pathway and promotes oncogenesis"

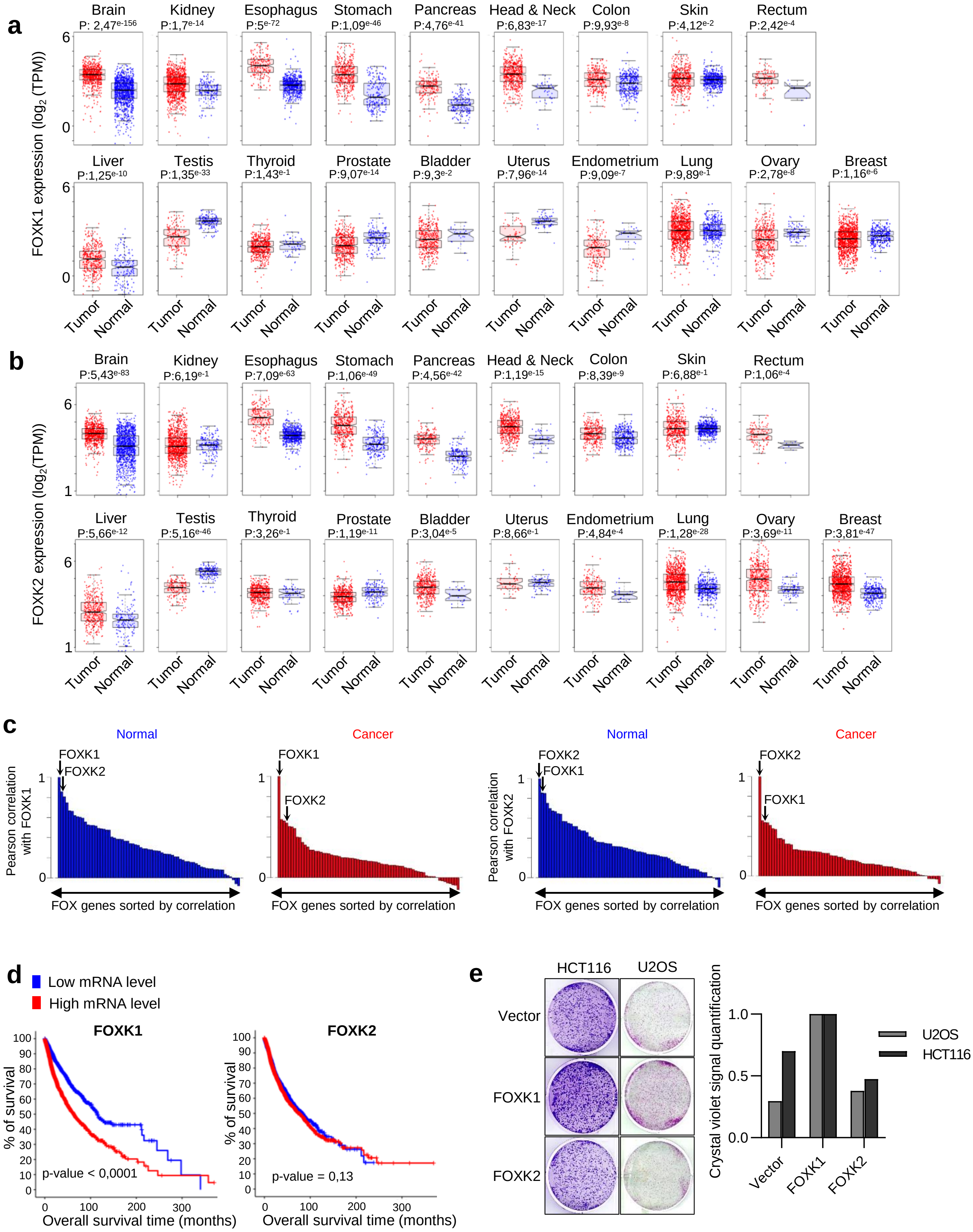


Extended Figure 1

**Extended Data Figure 1**:

**FOXK1 overexpression in cancer is associated with a poor prognosis**

**a**-**b**) Comparison of FOXK1 and FOXK2 expression between cancer and normal tissues. Cancer data were retrieved from TCGA TARGET GTEx dataset. P-value is calculated by Wilcoxson test. **c**) Comparison of co-expression of FOXK1, FOXK2 and other FOX genes between cancer and normal tissues. FOX genes were sorted by Pearson correlation with FOXK1 and FOKX2. **d**) Kaplan-Meier survival curve of patients from the TCGA database presenting high or low mRNA levels of FOXK1 or FOXK2 in cancer tissues. **e**) Proliferation of U2OS and HCT116 cells stably expressing FOXK1, FOXK2 or empty vector was analyzed by colony forming assay (CFA). Crystal violet signal for each condition was quantified using ImageJ and plotted in the right. Results from one representative experiment are shown.

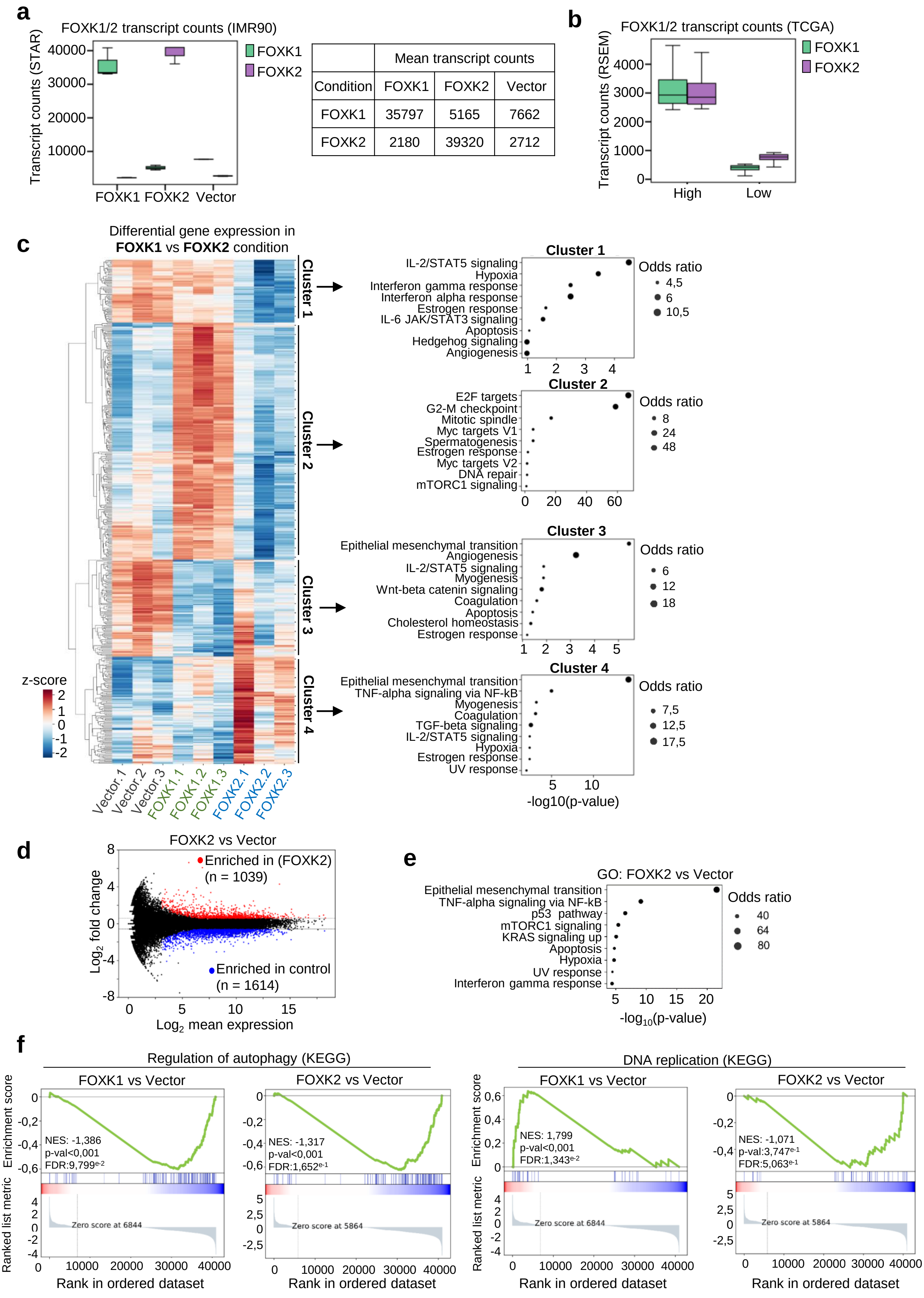


**Extended Figure 2**

**Extended Data Figure 2**:

**FOXK1 and FOXK2 regulate overlapping and specific gene expression programs**

**a**) Transcript counts of FOXK1 and FOXK2 from our RNA-seq experiment in IMR90 cells expressing either empty vector, FOXK1 or FOXK2. Mean transcript counts for each condition are represented in the adjacent table. **b**) Transcript counts from TCGA cancer samples were retrieved and categorized into two groups: the top 10% with the highest expression (mean FOXK1 = 3193, FOXK2 = 3115) and the top 10% with the lowest expression (mean FOXK1 = 398, FOXK2 = 757) of FOXK1 and FOXK2 transcripts. Counts for FOXK1 and FOXK2 in both group were plotted as boxplots. **c**) Heat map representing the transcript count (z-score) of genes differentially expressed between FOXK1 and FOXK2. GO analysis (MSigDB hallmark) was performed for each gene cluster. **d**) MA Plot representing the mean expression against the log fold change of genes when comparing IMR90 cells overexpressing FOXK2 with cells expressing the empty vector. **e**) GO analysis performed on genes differentially regulated between FOXK2 and control conditions. **f**) GSEA performed on genes deregulated (log fold change greater than 0.6) between conditions of FOXK1 overexpression and empty vector or FOXK2 overexpression and empty vector. Enrichment of genes associated to autophagy and DNA replication are represented.

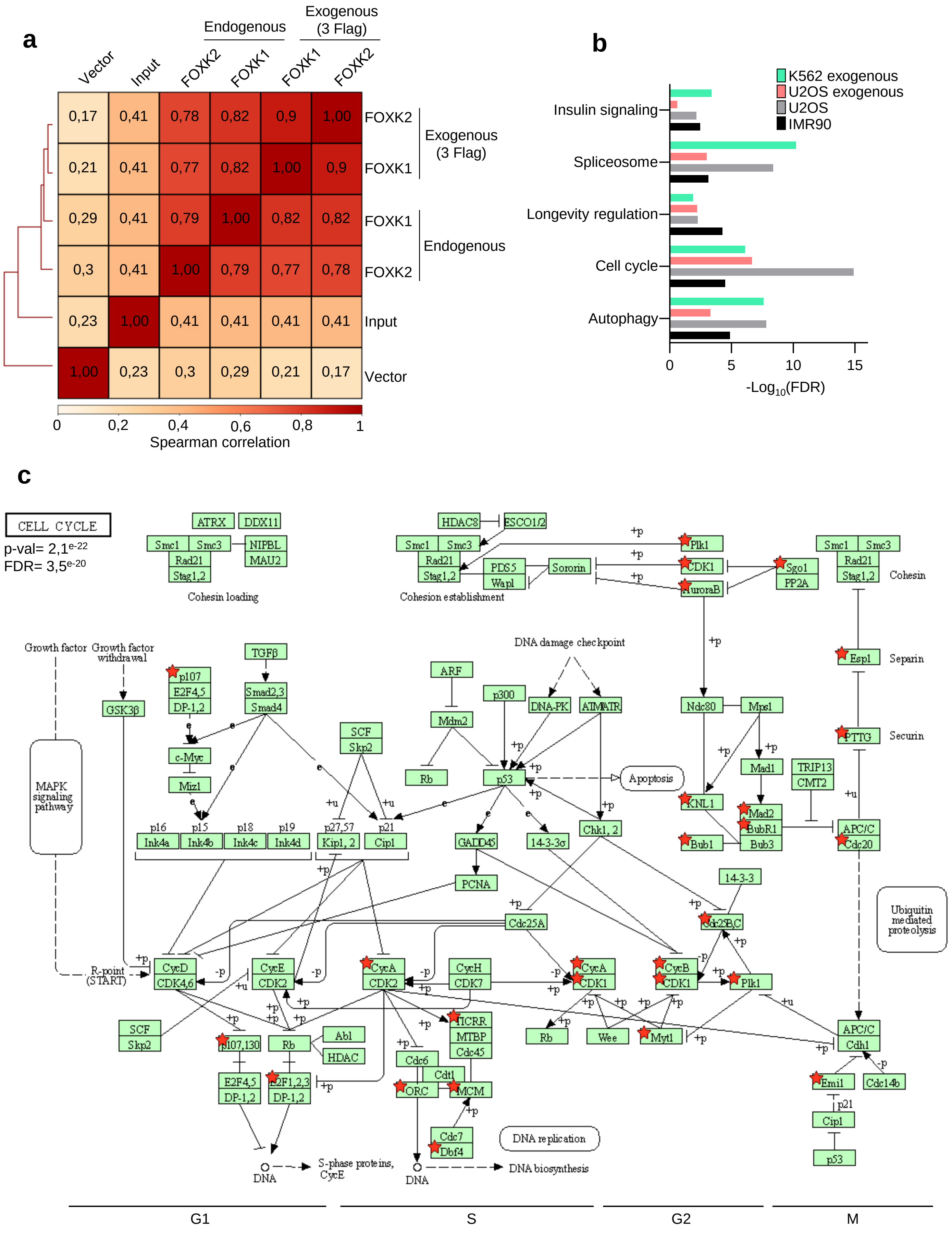


**Extended Figure 3**

**Extended Data Figure 3**:

**FOXK1 genome occupancy of cell cycle genes**

**a**) Correlation plot between endogenous and exogenous (Flag-tagged) ChIP-seq peaks of FOXK1 and FOXK2 in IMR90 cells. **b**) GO analysis of common promoters targeted by endogenous or exogenous FOXK1 in IMR90, U2OS and K562 as determined by ChIP-seq. **c**) GO analysis of genes up-regulated by FOXK1 in IMR90 found in the cell cycle pathway (KEGG) containing a ChIP-seq signal of FOXK1 in their promoters (red star).

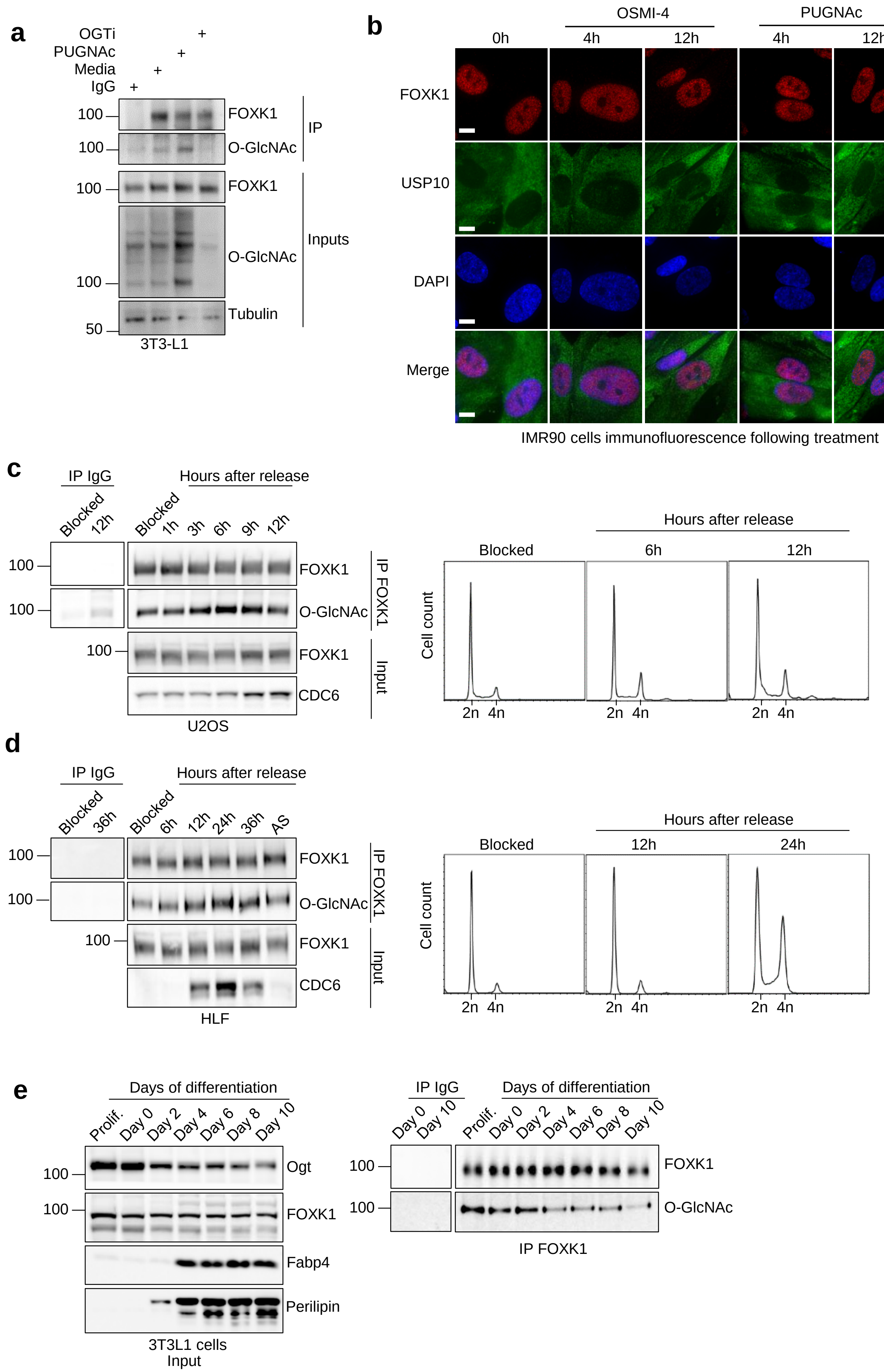


**Extended Figure 4**

**Extended Data Figure 4**:

**FOXK1 O-GlcNAcylation during cell cycle progression and cell differentiation**

**a**) Immunoprecipitation of endogenous FOXK1 and analysis of its O-GlcNAc levels in murine 3T3L1 cells treated with the OGA inhibitor, PUGNAc, or OGT inhibitor (OGTi). **b**) FOXK1 cellular localization following treatment with OGA inhibitor, OSMI-4, or with OGT inhibitor, PUGNAc, was analyzed by immunofluorescence in IMR90 cells. The non-relevant USP10 protein serves as a control for the cytoplasmic compartment. Representative of three independent experiments. **c**) U2OS cells were deprived of serum for 24h to synchronize cells in G1 phase. Cells were then stimulated with the addition of serum and FOXK1 was immunoprecipitated at different times to analyze its O-GlcNAcylation levels. CDC6 was used as a control of cell synchronization. **d**) Human lung fibroblasts (HLF) were synchronized by contact inhibition for several days to induce G0 entry. Cell cycle block release was performed by trypsinization and plating at low density in fresh medium. FOXK1 was immunoprecipitated at different times to analyze its O-GlcNAcylation. CDC6 was used as a control of synchronization. **e**) Pre-adipocytes 3T3-L1 were differentiated into adipocytes and FOXK1 was immunoprecipitated to analyze its O-GlcNAcylation levels upon differentiation. Fabp4 and Perilipin are markers of adipocytes.

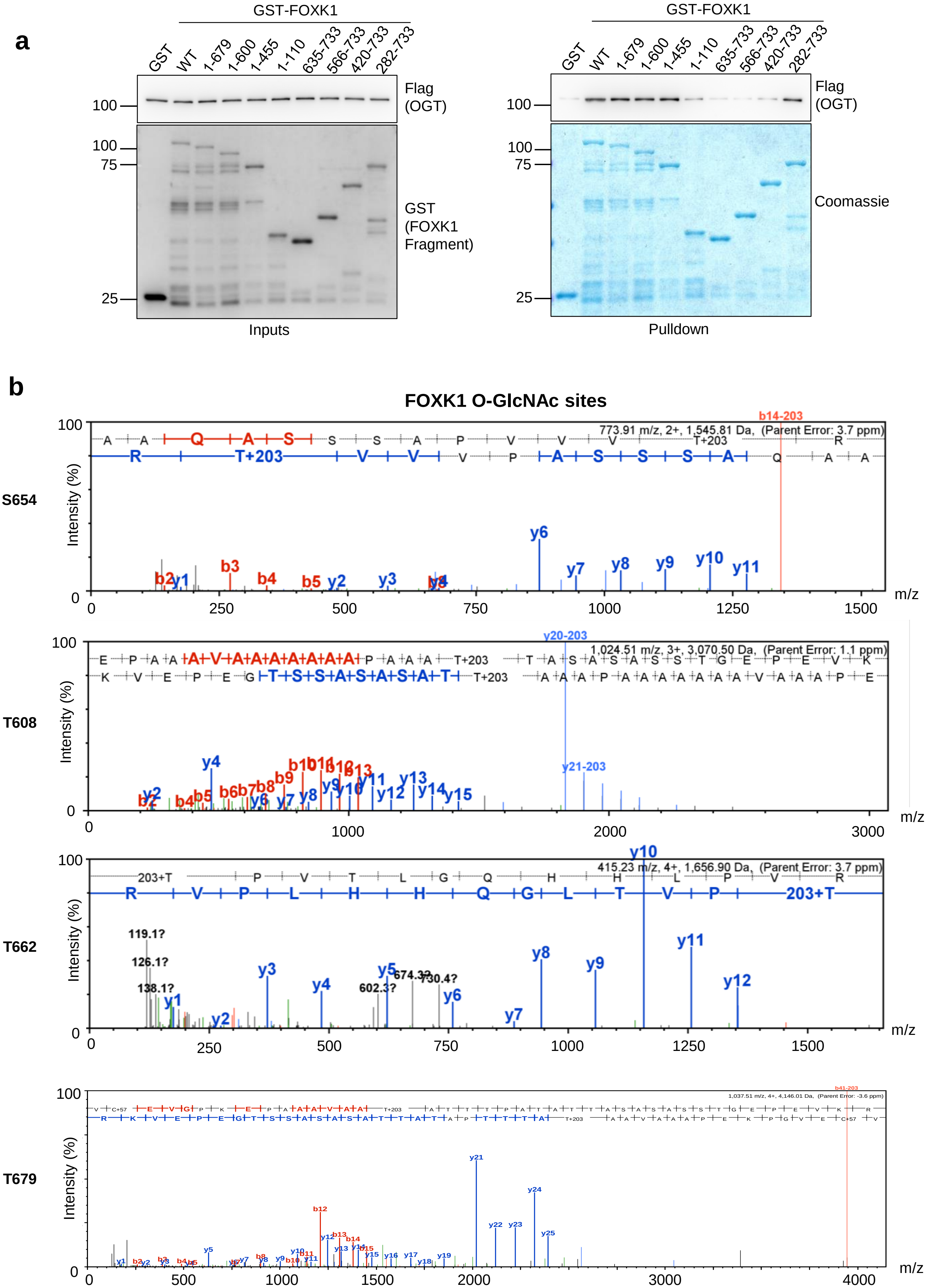


Extended Figure 5

### FOXK1 O-GlcNAc sites (continued)

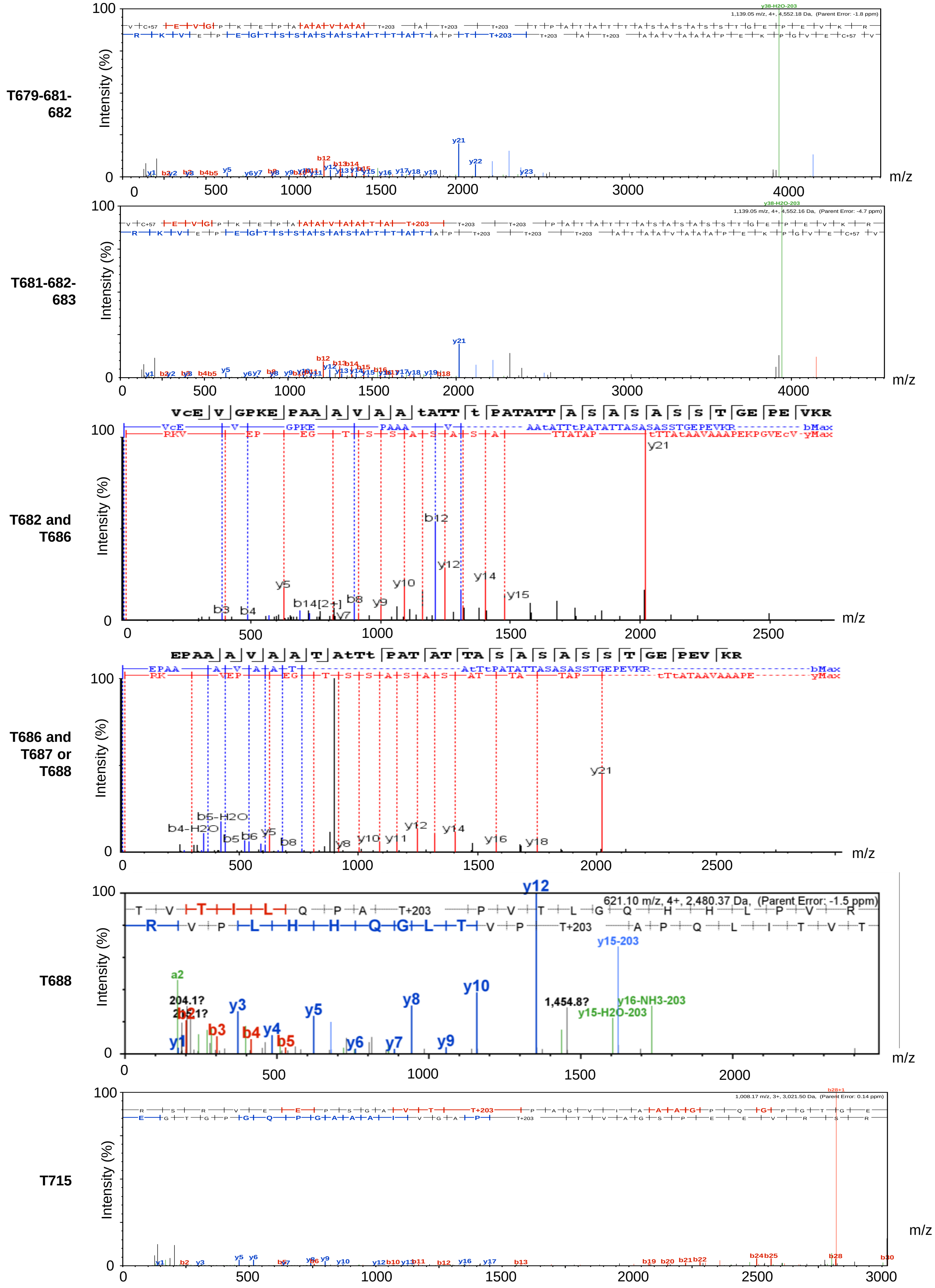


Extended Figure 5 (continued)

**Extended Data Figure 5**:

**Mapping of FOXK1 region and sites targeted by O-GlcNAcylation**

**A**) Recombinant GST-FOXK1 fragments pulldown with OGT to determine its interaction motif with FOXK1. Representative of two experiments. **B**) Mass spectra of FOXK1 residues targeted by O-GlcNAcylation.

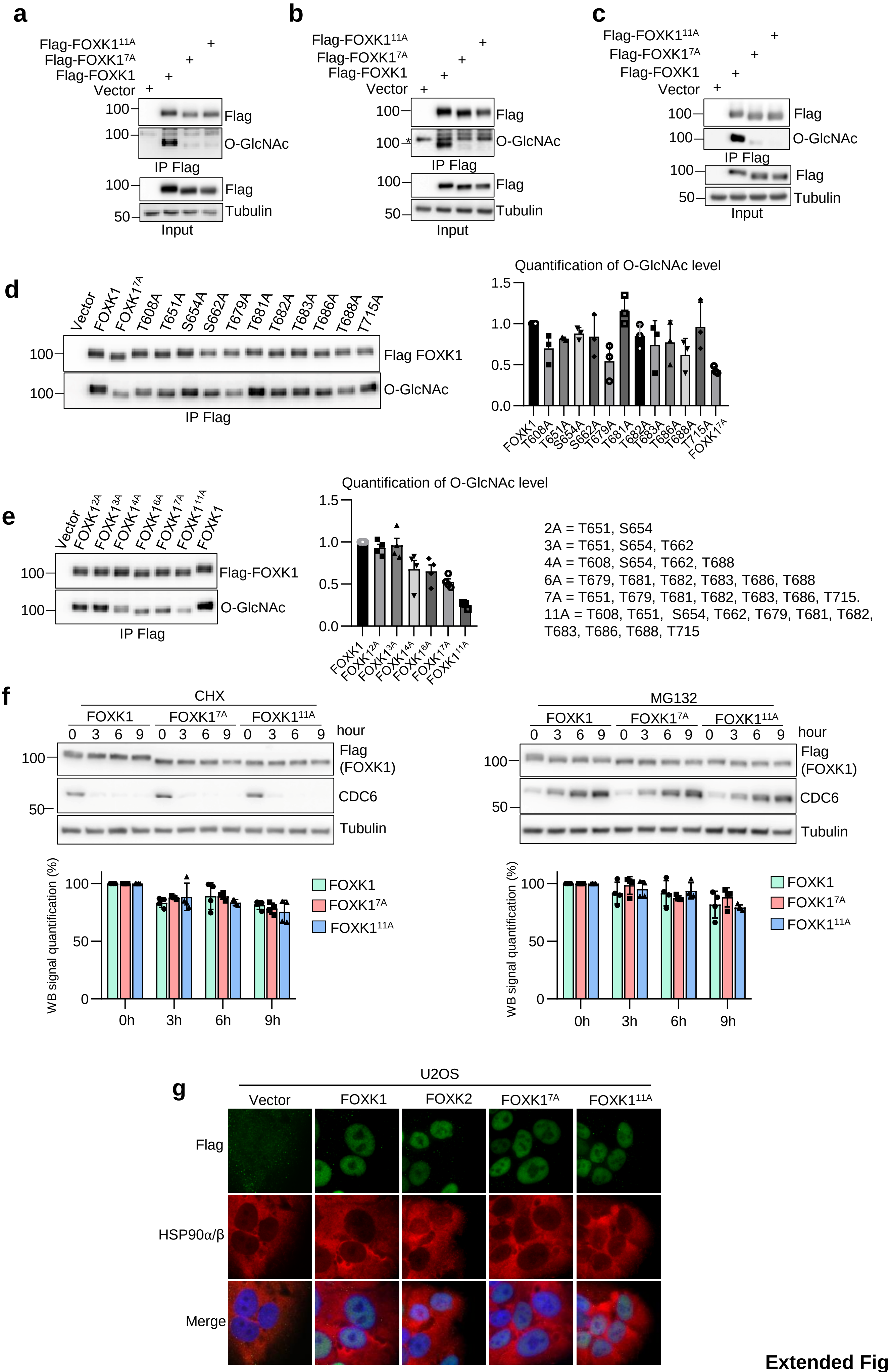


**Extended Figure 6**

**Extended Data Figure 6**:

**Characterization of FOXK1 O-GlcNAcylation sites and impact of O-GlcNAcylation on protein stability and localization.**

**a**) PC3 **b**) U2OS and **c**) K562 cells stably expressing Flag tagged version of FOXK1, FOXK1$^{7A}$ or FOXK1$^{11A}$ were harvested for Flag immunoprecipitation and O-GlcNAcylation detection. The star (*) represent non-specific bands in O-GlcNAc signal from U2OS cells. **d**) Transient transfection in HeLa cells of individual O-GlcNAc-modified residues mutation in FOXK1 (Flag tagged) to assess their O-GlcNAcylation levels after immunoprecipitation. The O-GlcNAc levels of FOXK1 mutants was quantified using ImageJ and plotted (right histogram) (n=3). **e**) Transient transfection in HeLa cells of combined O-GlcNAc-modified residues mutation in FOXK1 to assess O-GlcNAcylation levels following immunoprecipitation. The O-GlcNAc levels of FOXK1 mutants was quantified using ImageJ and plotted (right histogram) (n=4). **f**) U2OS cells overexpressing FOXK1, FOXK1$^{7A}$ or FOXK1$^{11A}$ were treated with 20 µg/ml cycloheximide or 20 µM MG132 and harvested for protein levels assessment by immunoblotting. CDC6 was used as a control for treatment efficacy. Flag-FOXK1 signal was quantified and normalized to tubulin signal. Representative of three independent experiments. **G**) Sub-cellular localization of exogenous FOXK1, FOXK1$^{7A}$, FOXK1$^{11A}$ and FOXK2 in U2OS cells. Detection of HSP90α/β was used as a control of the cytoplasm compartment. Representative of three independent experiments.

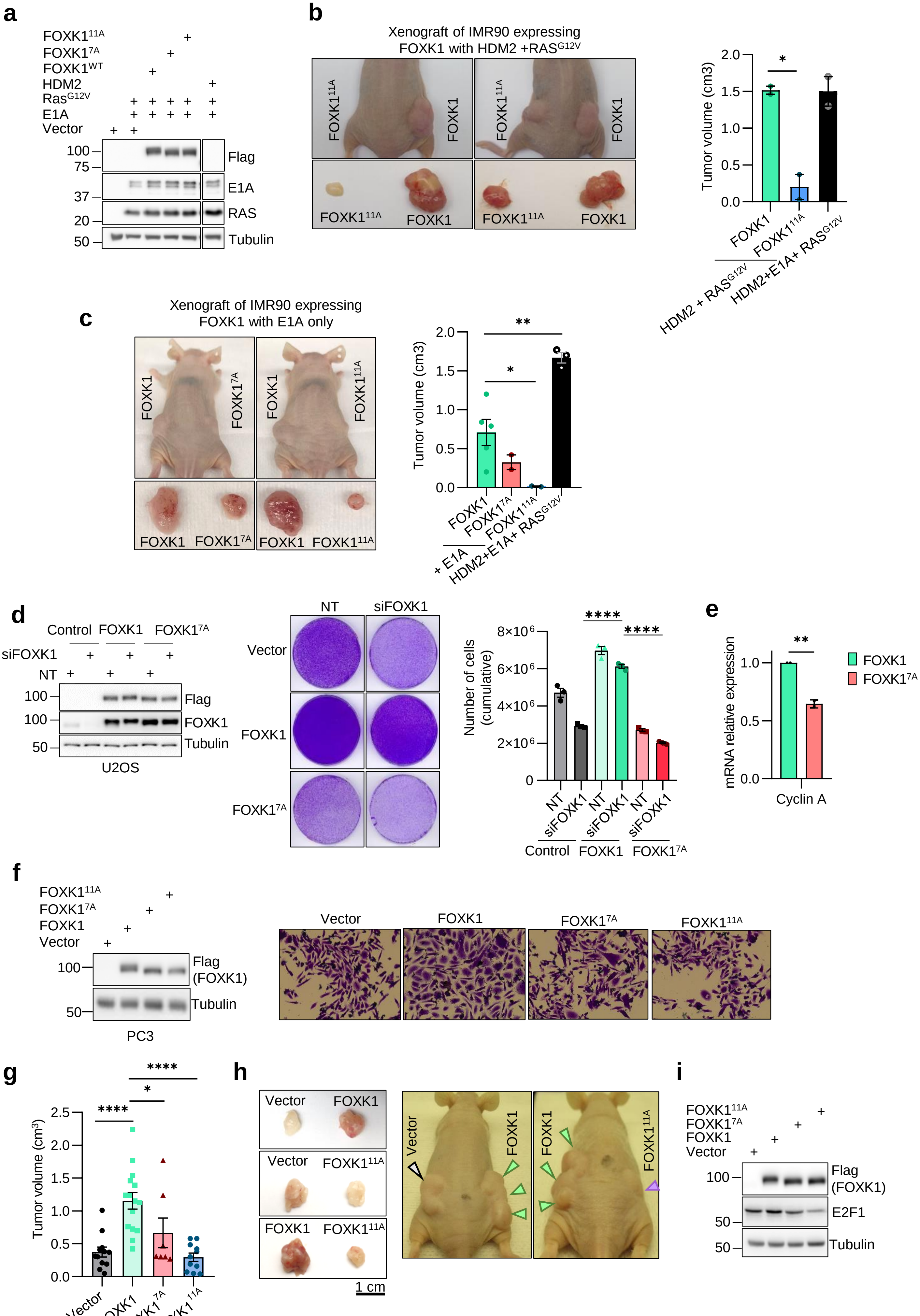


**Extended Figure 7**

**Extended Data Figure 7**:

**Effect of FOXK1 O-GlcNAcylation on tumor formation and progression**

**a**) Western blot of IMR90 expressing empty vector, FOXK1, $FOXK1^{7A}$ or $FOXK1^{11A}$ in combination with E1A and $RAS^{G12V}$. The full combination containing HDM2 + E1A + $RAS^{G12V}$ is also shown. **b**) Left: IMR90 tumors expressing HDM2 + $RAS^{G12V}$ in combination with FOXK1 or $FOXK1^{11A}$ at the time of harvest. Right: Graph representing final volume of tumors expressing FOXK1 or $FOXK1^{11A}$ in combination with HDM2 + $RAS^{G12V}$, and in comparison with tumors expressing HDM2 + E1A + $RAS^{G12V}$. **c**) Left: IMR90 tumors expressing FOXK1, $FOXK1^{7A}$ or $FOXK1^{11A}$ with E1A (minimal combination) at the time of harvest. Right: Graph of final size of tumors expressing FOXK1, $FOXK1^{7A}$ or $FOXK1^{11A}$ in combination with E1A, and in comparison with tumors expressing HDM2 + E1A + $RAS^{G12V}$. **d**) U2OS cells stably expressing empty vector, siRNA-resistant FOXK1 cDNA (FOXK1 or $FOXK1^{7A}$) were transfected with siRNA non-target (NT) or siRNA targeting endogenous FOXK1. Left: western-blotting depicting FOXK1 or $FOXK1^{7A}$ expression and the efficiency of endogenous FOXK1 depletion by siRNA. Middle: Cells were plated to perform colony forming ability (CFA). Right: Violet crystal was extracted from cells and intensity was quantified by absorbance (technical triplicates). **e**) mRNA expression of Cyclin A in U2OS expressing FOXK1 or $FOXK1^{7A}$ . Biological replicates. **f**) PC3 stably expressing empty vector, FOXK1, $FOXK1^{7A}$ or $FOXK1^{11A}$ were plated at low density for several days. Cells were stained with crystal violet. **g**) PC3 expressing empty vector, FOXK1, $FOXK1^{7A}$ or $FOXK1^{11A}$ were engrafted subcutaneously in the flanks of nude mice. Mice were sacrificed once the tumors reached the limit point. **h**) Images of tumors at the time of harvest. **i**) E2F1 protein levels were analyzed by western blotting on cell extracts of PC3 expressing empty vector, FOXK1, $FOXK1^{7A}$ or $FOXK1^{11A}$. Data are represented as mean ± SEM. One-way ANOVA with Dunnett's multiple comparisons was used (**b**, **c**, **d**, **g**). Statistical t-test (**e**) $*P < 0.05$, $**P < 0.01$, $****P < 0.0001$.

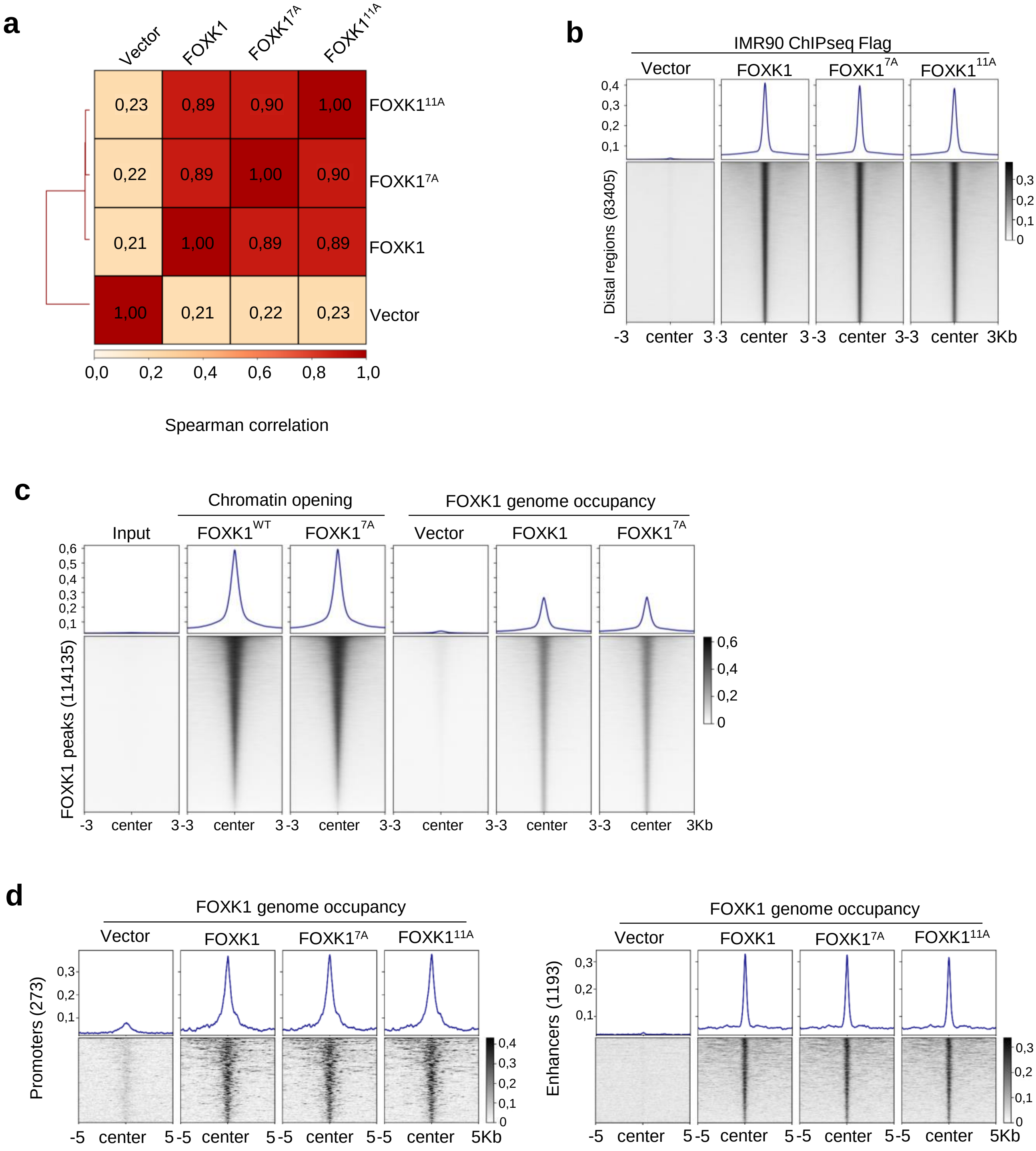


**Extended Figure 8**

**Extended Data Figure 8**:

**Effect of FOXK1 O-GlcNAcylation on genomic FOXK1 and BAP1 occupancy**

**a**) Correlation plot between exogenous ChIP-seq Flag signal for FOXK1, $FOXK1^{7A}$ and $FOXK1^{11A}$ in IMR90 cells. **b**) Chromatin occupancy of exogenous Flag tagged FOXK1, $FOXK1^{7A}$ and $FOXK1^{11A}$ in IMR90 cells on distal regions. Distal regions, corresponding to regions containing FOXK1 binding 1kb away upstream and downstream from TSS. **c**) Co-localization between FOXK1 and $FOXK1^{7A}$ ChIP-seq with opened chromatin regions from ATAC-seq experiments performed in U2OS cells overexpressing siRNA resistant cDNA of FOXK1 or $FOXK1^{7A}$. U2OS cells were treated with siRNA targeting FOXK1 for 72h before performing ATAC-seq experiment. ChIP-seq and ATAC-seq signals are centered on regions containing FOXK1 peaks. **d**) Occupancy of FOXK1, $FOXK1^{7A}$ and $FOXK1^{11A}$ assessed by ChIP-seq of 3-Flag tagged proteins in IMR90 on promoters and surrounding distal regions of genes whose expression is associated with FOXK1 overexpression.

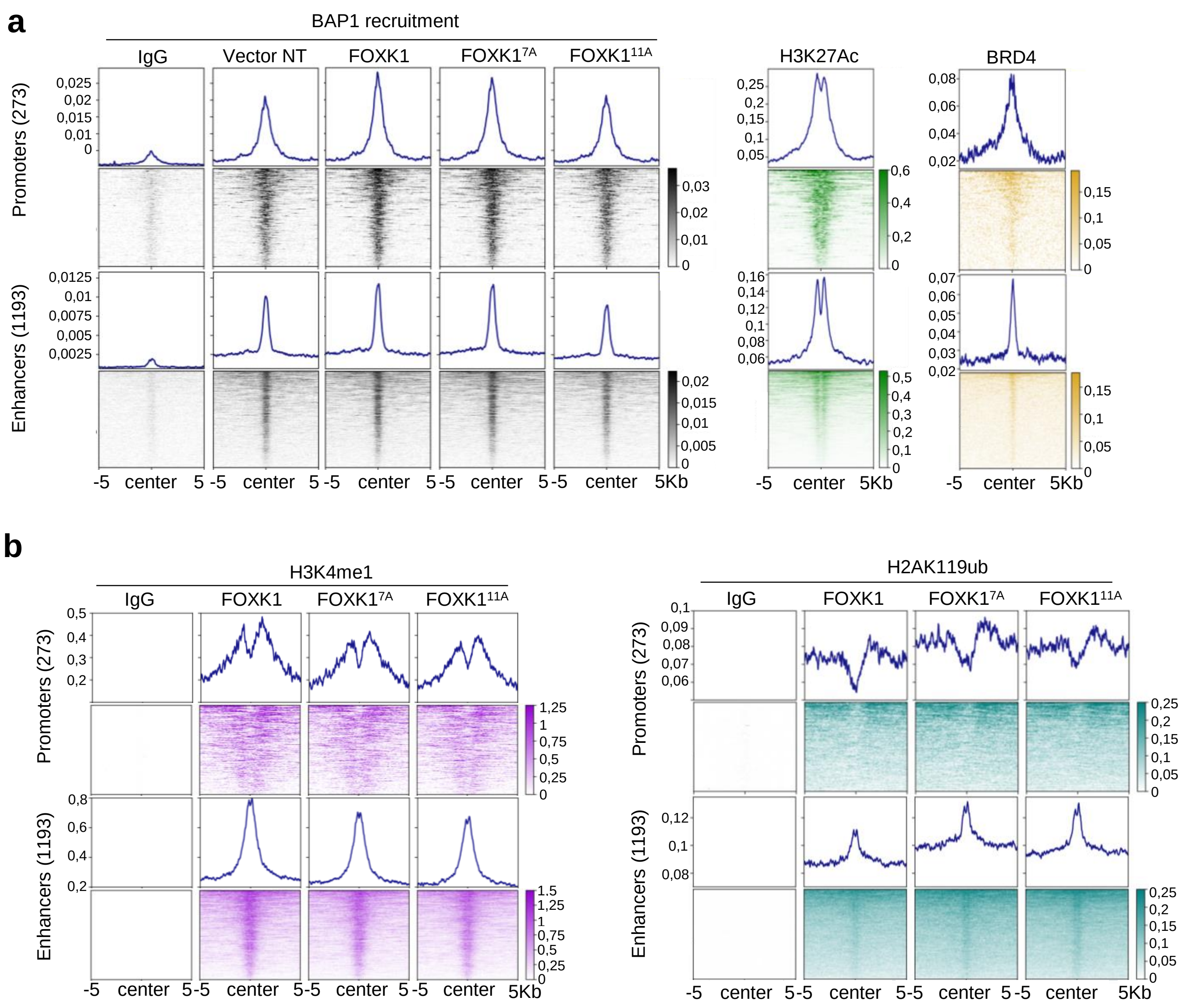


**Extended Figure 9**

**Extended Data Figure 9**:

**Effect of FOXK1 O-GlcNAcylation on epigenomic histone marks**

**a**) Analysis of BAP1 recruitment by Cut&Run in IMR90 cells expressing FOXK1, FOXK17A or FOXK111A on promoters of genes identified by RNA-seq as being differentially regulated by FOXK1 overexpression compared to FOXK2 or empty vector. Distal regions correspond to regions surrounding promoters at a distance greater than 1kb. These regions are enriched for H3K27Ac and BRD4 and were qualified as enhancers. **b**) Differential enrichment of H3K4me1 and H2AK119ub histone marks in IMR90 cells expressing FOXK1, FOXK17A or FOXK111A on the same promoters and distal regions.
